# Static growth alters PrrF- and 2-alkyl-4(1*H*)-quinolone regulation of virulence trait expression in *Pseudomonas aeruginosa*

**DOI:** 10.1101/765115

**Authors:** Luke K. Brewer, Weiliang Huang, Brandy Hackert, Maureen A. Kane, Amanda G. Oglesby

## Abstract

*Pseudomonas aeruginosa* is an opportunistic pathogen that is frequently associated with both acute and chronic infections, the latter of which are often polymicrobial. *P. aeruginosa* possesses a complex regulatory network that modulates nutrient acquisition and virulence, but our knowledge of these networks is largely based on studies with shaking cultures, which are not likely representative of conditions during infection. Here, we provide proteomic, metabolic, and genetic evidence that regulation by iron, a critical metallo-nutrient, is altered in static *P. aeruginosa* cultures. We identified type VI secretion as a target of iron regulation in *P. aeruginosa* in static but not shaking conditions, and we present evidence that this regulation occurs via PrrF sRNA-dependent production of 2-alkyl-4(1*H*)-quinolone metabolites. We further discovered that iron-regulated interactions between *P. aeruginosa* and a Gram-positive opportunistic pathogen, *Staphylococcus aureus*, are mediated by distinct factors in shaking versus static bacterial cultures. These results yield new bacterial iron regulation paradigms and highlight the need to re-define iron homeostasis in static microbial communities.

## INTRODUCTION

Antimicrobial-resistant bacterial pathogens represent a substantial health risk, raising concerns of a return to a “pre-antibiotic era.” One prominent example is multi-drug resistant (MDR) *Pseudomonas aeruginosa*, a Gram-negative opportunistic pathogen that is capable of establishing acute infections of the blood and lung in immunocompromised individuals. *P. aeruginosa* also causes chronic infections in surgical wounds, diabetic foot ulcers, and in the lungs of individuals with cystic fibrosis (CF) and chronic obstructive pulmonary disease (COPD). *P. aeruginosa* is the leading cause of morbidity among nosocomial pathogens, and is responsible for over 10% of all hospital-acquired infections (1-7). Therapeutic options for *P. aeruginosa* are rapidly dwindling due to the spread of antimicrobial resistance, underscoring the need for novel antimicrobials that can be used to treat MDR infections. To that end, a more complete understanding of the metabolic processes that contribute to *P. aeruginosa* survival and pathogenesis during infection is necessary for the discovery and development of effective therapeutics.

Iron is a required metallo-nutrient for most living organisms and has a significant impact on the establishment and progression of *P. aeruginosa* infections (8, 9). During infection, the host sequesters iron from invading pathogens through a process referred to as nutritional immunity (10-12). In response to iron starvation, *P. aeruginosa* expresses several virulence factors that cause host cell damage, presumably releasing iron and other nutrients from host cells (13-17). Iron starvation also induces the expression of multiple systems that mediate high affinity uptake of iron and heme (18, 19). Additionally, iron starvation induces expression of the PrrF small regulatory RNAs (sRNAs), which post-transcriptionally reduce expression of non-essential iron-dependent metabolic enzymes. The PrrF sRNAs are required for virulence in an acute murine lung infection model and are also highly expressed in the lungs of CF patients (9, 20, 21). However, the specific mechanisms by which the PrrF sRNAs contribute to mammalian infection remain unclear.

One prominent function of the PrrF sRNAs is their ability to indirectly promote the production of several small secreted metabolites collectively referred to as 2-alkyl-4(1*H*)-quinolones (AQs), which mediate a range of toxic activities against the host, as well as microbial pathogens that co-colonize *P. aeruginosa*-infected hosts. (20, 22-24). 2-alkyl-4-quinolone N-oxides (AQNOs) are potent cytochrome inhibitors that obstruct respiratory metabolism in mammalian and microbial cells (25-27). 2-alkyl-3-hydroxy-quinolone, which is more commonly referred to as PQS, and 2-alkyl-4-hydroxyquinolones (AHQs) both function as quorum signaling molecules and induce the expression of secreted factors that further inhibit growth of mammalian and microbial cells (28-31). AQ synthesis is initiated by enzymes encoded by the *pqsABCDE* operon, with PqsA catalyzing the first step with the conversion of anthranilate into anthraniloyl-CoA (32). The PrrF sRNAs promote AQ production by repressing the expression of the anthranilate degradation enzyme complexes AntABC and CatBCA (16), which degrade anthranilate to metabolites that enter the tricarboxylic acid (TCA) cycle. Thus, the PrrF sRNAs spare anthranilate for AQ production and are therefore required for optimal AQ production in iron-depleted conditions (9, 20, 23, 33).

In accordance with PrrF promoting AQ production in iron-depleted environments, we previously discovered that iron starvation enhances AQ-dependent antimicrobial activity against the Gram-positive pathogen *Staphylococcus aureus*, which can co-colonize with *P. aeruginosa* in chronic wounds and in CF lung infections (20, 23). Surprisingly, these previous studies showed that the *prrF* locus was not required for iron-regulated antimicrobial activity, and several possible explanations for this have been considered. One explanation that has not been investigated is that earlier studies used either agar plates or static liquid cultures, which are likely distinct from the shaking liquid cultures that have historically been used to study iron regulation. Whereas aerobic shaking cultures are exposed to constant aeration and perturbation to promote homogenous cultures, *P. aeruginosa* infections are typically more heterogeneous and feature planktonic as well as adherent biofilm communities of *P. aeruginosa*, particularly during chronic infections. Changes in the iron metabolism and iron regulatory factors in response to aeration and perturbation have been similarly documented in other bacterial species, suggesting a regulatory link between iron homeostasis mechanisms and static growth that may be common among other bacterial species (34, 35). Because of the broad impact of iron on *P. aeruginosa* physiology, we hypothesized that static growth would likewise alter the iron-regulated metabolism in *P. aeruginosa*, potentially revealing novel iron regulatory pathways that are more relevant during infection.

In the current study, we sought to identify targets of iron and PrrF regulation during static growth of *P. aeruginosa* using an unbiased proteomics-based approach. Our results revealed a number of proteins that were differentially affected by iron and PrrF in static versus shaking cultures. Amongst these were proteins for phenazine biosynthesis, which were repressed by iron in static conditions, a result that is in contrast to our own shaking cultures and recent studies showing that phenazine synthesis genes are iron-induced in shaking conditions (12, 36). We also discovered PrrF positively affects the expression of the Hcp1 secretion island II type VI secretion system (HSI-II T6SS) in *P. aeruginosa*, and we provide evidence that this regulation is due to PrrF-regulated AQ production. Moreover, while PrrF is not required for antimicrobial activity against *S. aureus* in static conditions, we found that it still promotes the production of AQs in static conditions. However, the relative abundances and roles of individual AQ species for interspecies interactions varied between shaking versus static cultures, thereby altering the requirement of PrrF for antimicrobial activity. Lastly, we show that reduced oxygen availability in static cultures is not solely responsible for the changes in iron regulation observed in this study. Combined, these results indicate that static bacterial communities use unique regulatory and metabolic networks to adapt to decreased nutrient availability, likely affecting the production of key virulence determinants.

## MATERIALS AND METHODS

### Bacterial strains and growth conditions

Bacterial strains used in this study are listed in **Supplementary Table S1**. Lysogeny broth (LB) and agar (LA) were prepared using 10g/L NaCl (Sigma, St Louis, MO), 10g/L tryptone, and 5g/L yeast extract, 15g/L agar when applicable (Becton-Dickinson, Franklin Lakes, NJ). *P. aeruginosa* and *S. aureus* strains were both routinely grown on LA from freezer stock. Five isolated colonies of each strain were selected from overnight-incubated agar plates and inoculated into 5mL LB. For iron starvation studies, bacterial strains were subcultured into Chelex-treated and dialyzed trypticase soy broth (DTSB) prepared as previously described (16). Media dialysis was carried out using Spectra/Por®2 dialysis membrane tubing (29mm diameter) with a molecular weight cutoff of 12-14kD (Repligen, Waltham, Ma).

For co-culture assays, quantitative real time PCR (qRT-PCR) analysis, and mass spectrometry-assisted metabolite and proteome studies, *S. aureus* and *P. aeruginosa* cultures were diluted to an absorbance (OD_600_) of 0.08 and 0.05, respectively, into 1.5mL of DTSB with (high iron) or without (low iron) 100 µM FeCl_3_ supplementation,. Mono- and co-cultures of *P. aeruginosa* and *S. aureus* were prepared in DTSB media and incubated at 37°C for 18 hours in a shaking incubator. Shaking aerobic cultures were incubated in 1.5mL of DTSB media in 14mL round bottom tubes and closed with a foam stopper to allow for sufficient aeration. Static cultures were incubated in 1.8mL DTSB media in 6-well polystyrene culture plates, which were covered in breathe-easy wrap to prevent evaporation. During incubation, cultures were grown at a shaking rate of either 250rpm (shaking conditions) or 0rpm (static conditions). Microaerobic co-cultures were grown in identical growth conditions as aerobic shaking cultures, except culture tubes were placed into an air-tight candle jar in the presence of a CampyPak (BD Diagnostics, NJ, USA). Sealed candle jars were secured in a shaking incubator and incubated for 18 hours at 37°C and a shaking rate of 250rpm.

### Quantitative label-free proteomics

Cell cultures were prepared for proteomics analysis as described previously (36). Briefly, cells were harvested by centrifugation and washed in phosphate-buffered saline prior to lysis in 4% sodium deoxycholate. Lysates were washed, reduced, alkylated and trypsinolyzed on filter (37, 38). Tryptic peptides were separated using a nanoACQUITY UPLC analytical column (BEH130 C18, 1.7 μm, 75 μm x 200 mm, Waters) over a 165-minute linear acetonitrile gradient (3 – 40%) with 0.1 % formic acid on a Waters nano-ACQUITY UPLC system and analyzed on a coupled Thermo Scientific Orbitrap Fusion Lumos Tribrid mass spectrometer. Full scans were acquired at a resolution of 120,000, and precursors were selected for fragmentation by higher-energy collisional dissociation (normalized collision energy at 32 %) for a maximum 3-second cycle. Tandem mass spectra were searched against *Pseudomonas* genome database PAO1 reference protein sequences (39) using Sequest HT algorithm and MS Amanda algorithm with a maximum precursor mass error tolerance of 10 ppm (40, 41). Carbamidomethylation of cysteine and deamidation of asparagine and glutamine were treated as static and dynamic modifications, respectively. Resulting hits were validated at a maximum false discovery rate (FDR) of 0.01 using a semi-supervised machine learning algorithm Percolator (42). Label-free quantifications were performed using Minora, an aligned AMRT (Accurate Mass and Retention Time) cluster quantification algorithm (Thermo Scientific, 2017). Protein abundance ratios between the high iron cultures and the low iron cultures were measured by comparing the MS1 peak volumes of peptide ions, whose identities were confirmed by MS2 sequencing as described above. Gene function and pathway analysis was completed using information from the *Pseudomonas* genome database (39), KEGG database (43), *Pseudomonas* metabolome database (44), and the STRING database (45).

### Real time PCR analysis

Quantitative real time PCR (qRT-PCR) analysis was conducted on at least three biological replicates of the PAO1 reference strain, the isogenic Δ*prrF* mutant, or Δ*pqsA* mutant grown in DTSB supplemented with (high iron) or without (low iron) 100 µM FeCl_3_. *P. aeruginosa* cells were lysed after 18 hours using 2.5mg/mL lysozyme and incubated at 37°C for 30 minutes. RNA extraction was performed using the RNeasy mini kit (QIAGEN, Germantown, MD). Real-time qualitative PCR analysis was performed as described previously (9) using an Applied Sciences StepOne Plus Real Time PCR System (Life Technologies, Carlsbad, CA). Primer and probe sequences used for gene expression analysis are listed in **Supplementary Table S2**. Quantitation of cDNA was carried out using standard curves generated by performing qPCR of individual gene targets using cDNA reverse-transcribed from serial dilutions of RNA. *P. aeruginosa* expression data was normalized to *oprF* expression in aerobic shaking conditions, or *omlA* in either microaerobic shaking conditions or static growth condition, as expression of these genes is not impacted by iron in each respective growth condition (9).

### Quantitation of AQs using liquid chromatography-tandem mass spectrometry

AQ quantitation was carried out as previously reported (46). Briefly, 300 L of culture were harvested and spiked with a 25 µM stock nalidixic acid internal standard to a final concentration of 500 nM, followed by an extraction using ethyl acetate with 0.1% acetic acid. The organic phase containing extracted AQs was harvested from each sample, dried down, and resuspended in 300 µL of 0.1% formic acid suspended in 1:1:1 (v/v/v) mixture of methanol, water, and acetonitrile. Extracts were analyzed via quantitative liquid chromatography-tandem mass spectrometry (LC-MS/MS) using multiple reaction monitoring (MRM) performed on a Waters ACQUITY UHPLC coupled to a Xevo TQ-XS tandem quadrupole mass spectrometer in positive ion mode.

### Quantification of viable cells in co-culture studies

400 µL of *P. aeruginosa* and *S. aureus* mono- and co-cultures were harvested at 15000 rcf for 5 minutes. Cell pellets were resuspended in 400 µL of 0.1% Triton-X in phosphate-buffered saline (PBS) and vortexed rigorously. From these resuspensions, serial dilutions were prepared in 0.1% Triton-X in PBS. 10 µL of each dilution was spotted onto Baird-Parker agar plates and Pseudomonas Isolation Agar (PIA) plates to select for *S. aureus* and *P. aeruginosa* growth, respectively. Upon spotting 10 µL of dilutions onto agar plates, the plates were tilted to facilitate even drips across the plate surface. PIA plates were incubated at 37°C for 18 hours and colony forming units (CFU) were counted. Baird-Parker plates were incubated for 37°C for 48 hours to allow for adequate growth of *S. aureus* small-colony variants observed in some assays prior to counting CFUs.

### Statistics

Statistically significant changes in cell viability, RNA gene expression, and AQ concentrations between treatment groups were identified using one-way ANOVA two-tailed on Prism 6, using either Tukey’s, Sidak’s, or Dunnett’s post test for multiple comparisons, with a significance threshold of *p* value < 0.05. Protein expressions that changed 2-fold or more with an FDR adjusted *p*-value < 0.05 were considered statistically significant.

## RESULTS

### Proteomics reveals altered iron-regulation in static *P. aeruginosa* cultures

Our current knowledge of *P. aeruginosa* iron regulation is largely restricted to shaking cultures (14, 16, 19, 23, 36, 47). To determine whether global iron regulatory pathways are altered in static cultures *of P. aeruginosa*, we applied a label-free, LC-MS/MS-based proteomics methodology recently described by our laboratories (36). Using this unbiased approach, we determined the proteomes of static and shaking cultures of PAO1 as well as the isogenic Δ*prrF* mutant grown in Chelexed and dialyzed TSB medium (DTSB), supplemented with or without 100 µM FeCl_3_. Mass spectra were identified using a PAO1 reference proteome, which were in turn validated to a false discovery rate (FDR) of 0.01 to ensure proper assignment of protein identities.

Initially, we sought to confirm whether iron regulatory effects previously identified in shaking conditions were similar to shaking cultures analyzed in the current study. For this, we identified proteins whose levels showed a statistically significant (FDR adjusted *p*-value <0.05) log fold change (LFC) > 1 upon iron starvation. This analysis revealed 178 and 191 proteins that were significantly induced and repressed, respectively, by iron starvation (**Supplementary Dataset 1**). As previously observed (14, 19, 48-52), iron starvation significantly increased levels of proteins for the siderophore (pyoverdine and pyochelin) and heme uptake systems, as well as several iron-regulated virulence factors, during shaking growth (**Fig. 1A**). Also as previously observed, iron-containing proteins involved in the tricarboxylic acid (TCA) cycle and oxidative metabolism, including aconitase A (AcnA), succinate dehydrogenase B (SdhB), and catalase (KatA), were induced by iron in a PrrF-dependent manner under shaking conditions (**Fig. 1B**) (16, 53). We also observed PrrF-dependent iron regulation of more recently identified PrrF targets in shaking conditions, including proteins involved in Fe-Sulfur cluster biogenesis (IscS, IscU) and amino acid metabolism (IlvD) (**Fig. 1B**) (36). Contrary to previous real-time PCR and microarray analyses showing iron induction of the *antABC* mRNAs (36), neither iron supplementation nor *prrF* deletion significantly affected levels of the AntABC proteins for anthranilate degradation under shaking conditions (**Fig. 1B**). Overall however, this analysis replicated many of the global iron regulatory pathways observed in previous studies of shaking *P. aeruginosa* cultures.

**Figure 1.**
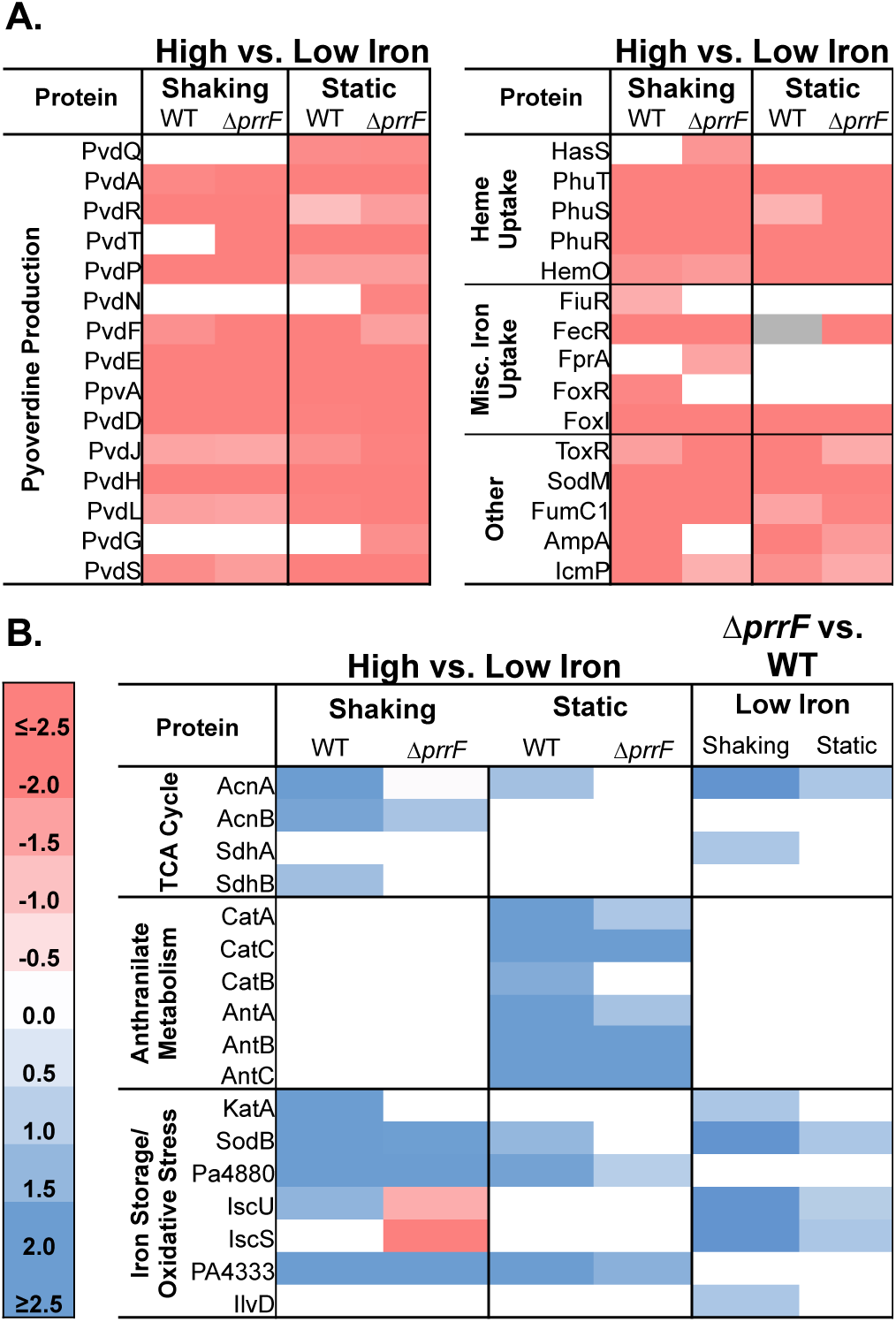
Static growth reduces the effects of classical PrrF regulation. Heatmaps showing the log_2_ fold change (LFC) of **(A)** known iron-repressed proteins and **(B)** known PrrF-regulated proteins, in the indicated strains grown in high versus low iron conditions or in the Δ*prrF* mutant versus wild type PAO1 grown in low iron conditions. Wild type PAO1 and Δ*prrF* strains were inoculated into 1.5mL DTSB supplemented with FeCl_3_ and incubated in shaking or static conditions as described in the Materials and Methods. LC-MS/MS-based proteomics analysis showed statistically significant protein regulation (FDR adjusted p-value < 0.05) of known iron and PrrF targets, as indicated by at least 2-fold induction or repression (i.e. 1 LFC) in response to treatment. Gray boxes indicate proteins that were undetected in one or more condition. White boxes indicate no significant change in gene expression was observed.

We next determined whether known targets of PrrF and iron regulation under shaking conditions were similarly regulated under static conditions. Iron repression of siderophore and heme uptake proteins was largely retained during static growth, indicating that these iron regulatory pathways are not altered under static conditions (**Fig. 1A**). In contrast, iron and PrrF regulation of several proteins involved in the TCA cycle or oxidative stress protection was reduced or eliminated in static conditions (**Fig. 1B**). We also noted robust iron induction of the anthranilate degradation proteins AntABC and CatBCA, which occurred in a PrrF-independent manner (**Fig. 1B**), in agreement with trends we observed in subsequent qRT-PCR analysis of *antR* and *antA* under identical growth conditions **(Fig. S1B**,**C)**. Combined, these data indicate that PrrF-dependent iron regulatory pathways are altered in static conditions.

To determine if there were additional iron or PrrF regulatory effects that were unique to either static or shaking growth, we mined our proteomics dataset for proteins whose levels were altered by iron depletion in either static or shaking conditions, but not in both, by an LFC ≥ 1 and an FDR adjusted *p* value ≤ 0.05. Approximately 410 proteins demonstrated significantly altered iron regulation in response to growth conditions. Of these, 126 of these were specifically iron induced and 93 were iron repressed in shaking but not static conditions, while 78 were iron induced and 129 were iron repressed only in static conditions (**Supplementary Table S3-7 and Dataset S1**). Approximately 50% of the proteins affected by iron supplementation in static conditions were unaffected by iron in the Δ*prrF* mutant (**Supplementary Table S3-7 and Dataset S1**), indicating PrrF still contributes to iron regulation during these growth conditions, despite the loss of PrrF-mediated iron regulation on previously identified targets as shown in **Fig. 1B**.

STRING network analysis was next used to identify relationships between the proteins within each of these four groups (45). Several of the proteins that were induced by iron in shaking conditions but not in static conditions were related to motility, including proteins for flagellar assembly (FlgA, FlgL, FliC, and FliI), a chemotaxis-associated protein (ChpA), and proteins involved in twitching pilus formation (PilV, PilY2, and FimU) **(Supplementary Dataset S1).** Proteins that were significantly induced by iron depletion in static but not shaking cultures included numerous enzymes required for synthesis of the redox cycling phenazine metabolites and the pyochelin siderophore, as well as proteins encoded by the HSI-II T6SS locus (**Fig 2A, Supplementary Fig S4-5)**. Iron regulation of the phenazine and pyochelin synthesis proteins occurred in a PrrF-independent manner, while iron regulation of many of the T6SS proteins was either reduced or lost in the Δ*prrF* mutant (**Fig. 2B**). Moreover, we found that *prrF* deletion had a negative effect on T6SS proteins in static but not shaking conditions **(Supplementary Dataset S1**). The identification of T6SS as iron- and PrrF-regulated during static but not shaking conditions was particularly intriguing, as this contact-dependent secretion system is known to mediate virulence and interbacterial interactions that may be more relevant in static conditions than in highly perturbed, shaking conditions. No complementarity was identified between the PrrF sRNAs and the mRNAs encoding these proteins, thus we hypothesized that PrrF-regulated expression of these proteins occurs via an indirect mechanism.

**Figure 2.**
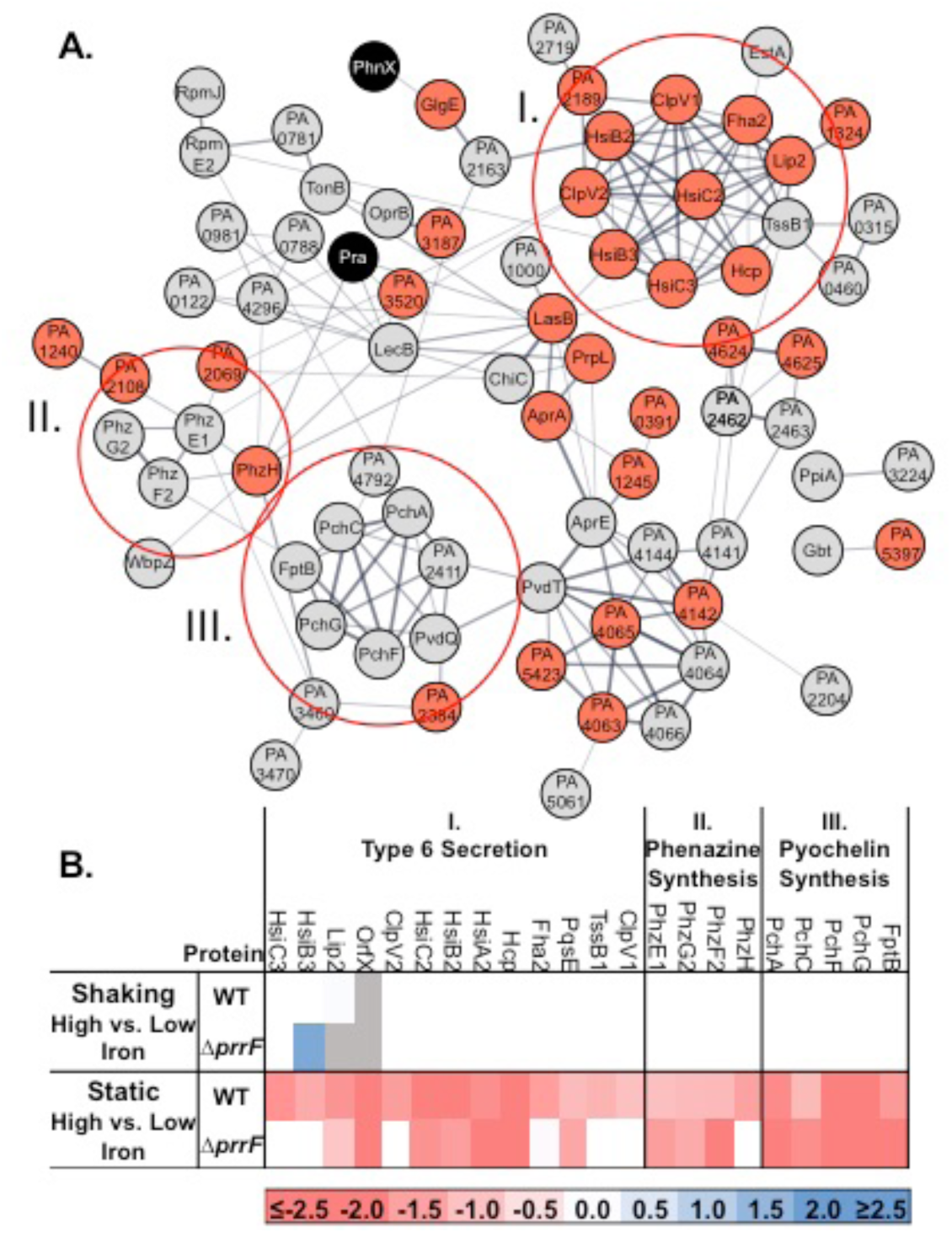
Proteomics reveals novel targets of iron regulation in static conditions. Wild type PAO1 and the Δ*prrF* strain were inoculated into 1.5mL DTSB media and supplemented with either low iron (0 µM) or high iron (100 µM) and incubated in either shaking or static conditions. Statistically significant iron regulation (FDR adjusted p-value < 0.05) in static and shaking conditions was identified by at least 2-fold (i.e.1 LFC) induction in response to iron starvation as described in Materials and Methods. **(A)** Network analysis of differentially regulated genes was carried out using STRING database software, which revealed distinct several virulence-associated proteins that were similarly impacted by iron starvation under static conditions. Line thickness indicates strength of data support for association between two proteins, as calculated by STRING network analysis (45). Red nodes represent proteins that were significantly regulated by PrrF in low iron static conditions, gray nodes indicated proteins that do not exhibit regulation by PrrF. Black nodes represent proteins that exhibited iron regulation in static conditions but were not detected in one or more iron conditions in shaking cultures. **(B)** Heatmaps of differentially regulated proteins encoded by the HSI-II T6SS, phenazine, and pyochelin biosynthetic operons. Gray boxes indicate proteins that were not detected in one or more condition. White boxes indicate that no significant change in protein levels was observed.

### HSI-II T6SS genes are differentially regulated by AQs in shaking versus static conditions

No previous reports have demonstrated a regulatory link between the PrrF sRNAs and T6SS, nor were we able to identify complementarity between PrrF and T6SS secretion mRNA sequences. Previous reports, however, have demonstrated that PrrF sRNAs are necessary for full AQ production via repression of the anthranilate-degrading enzymes (9, 33). AQs, in turn, induce expression of several virulence-associated genes, including genes involved in T6SS (54, 55). We therefore sought to determine whether reduced levels of T6SS proteins in the Δ*prrF* mutant during our proteomics analysis were due to a partial loss of AQ production.

To investigate this idea, we first performed targeted expression analysis of T6SS genes in our wild type, Δ*prrF*, and Δ*pqsA* strains grown in static conditions using quantitative real time PCR (qRT-PCR). Specifically, we analyzed the expression of genes encoding proteins that demonstrated PrrF dependent iron regulation in our proteomics analysis, including *clpV2, fha2*, and *lip2*. In agreement with our proteomics analysis, the Δ*prrF* mutant exhibited statistically significant decreases in expression of each of these genes when grown in static conditions (**Fig. 3A**). The Δ*pqsA* mutant demonstrated even more dramatic decreases in the expression of *clpV2, fha2*, and *lip2* relative to wild type PAO1 (**Fig. 3A**), supporting the notion that AQs are necessary for the expression of T6SS genes in static cultures. Similar AQ-dependent expression of other T6SS genes was also observed (**Fig. S2).** In contrast, neither *prrF* nor *pqsA* were required for T6SS expression in shaking conditions (**Fig. 3B and Fig. S2**), supporting the hypothesis that AQ regulation of T6SS is enhanced by static growth. In agreement with our proteomics, T6SS expression was iron regulated in static cultures of wild type PAO1, but not in shaking cultures (**Fig. 3A,B)**, with expression of *clpV2, fha2*, and *lip2* repressed by an average of 4.07-, 3.12-, and 3.86-fold in high iron static cultures, respectively. Iron repression of these genes was diminished or was not statistically significant in static Δ*pqsA* cultures (*clpv2* repressed 2.40-fold; *fha2* repressed 1.72-fold; *lip2* repressed 2.25-fold). Interestingly, expression of T6SS synthesis proteins was still repressed by iron in static Δ*prrF* cultures, with expression of *clpV2, fha2, and lip2* repressed by an average of 3.78-fold, 3.63-fold, and 3.83-fold in high iron static conditions, respectively, suggesting that PrrF is not the sole mediator of iron-regulated T6SS gene expression in PAO1. We noted that expression of the T6SS genes was greatly reduced in low iron shaking cultures of PAO1, with C_t_ values 2-3 higher than what was observed in low iron static cultures of PAO1 (data not shown). Thus, shaking growth appears to reduce T6SS expression by PAO1 when grown in DTSB low iron medium, masking iron, PrrF, and AQ regulation to be observed.

**Figure 3.**
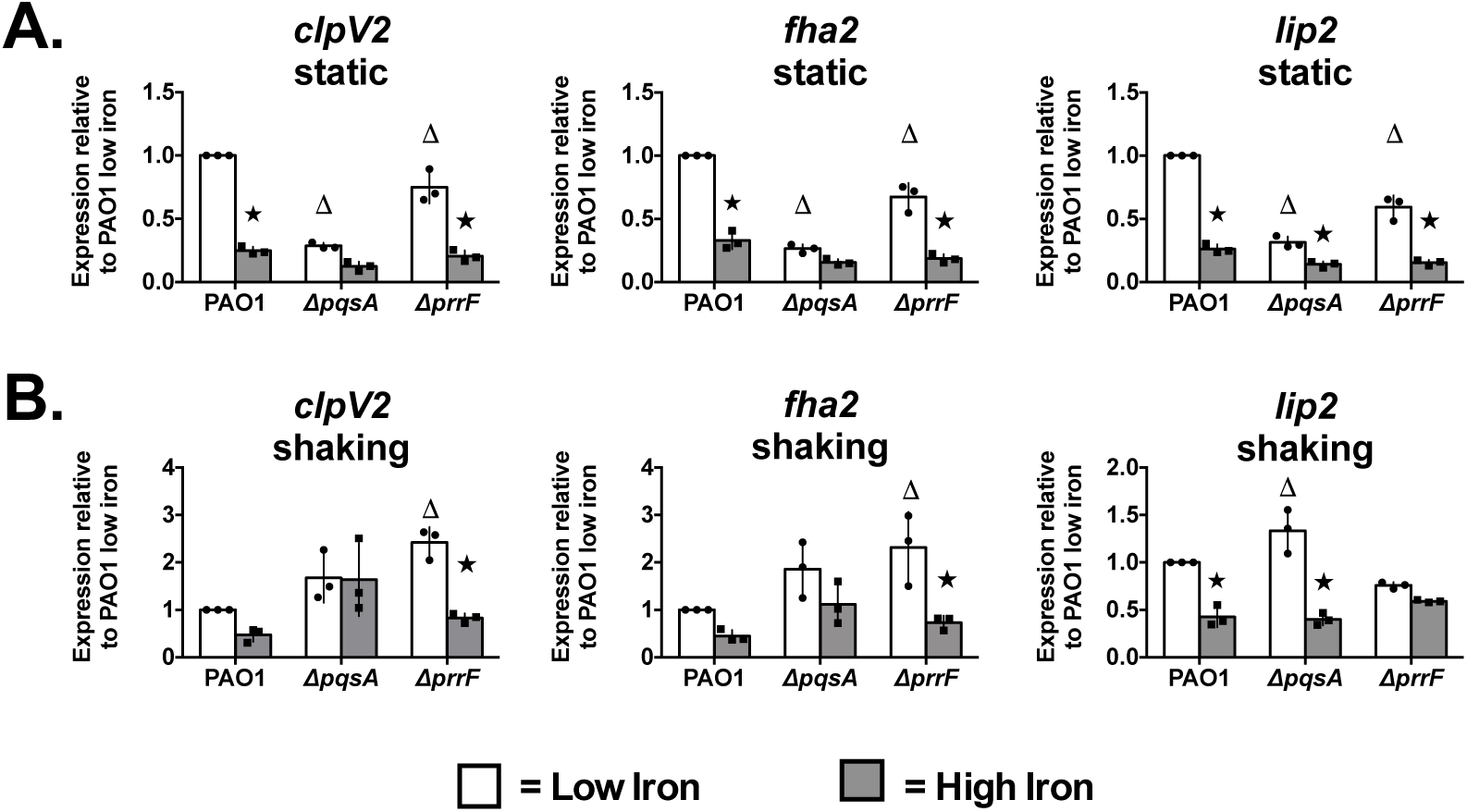
PrrF and AQs differentially regulate expression of T6SS genes in static and shaking conditions. Cultures were grown in 1.5 mL of DTSB media supplemented with either 0 µM FeCl_3_ or 100 µM FeCl_3_ in 14mL polystyrene round bottom culture tubes for 18 hours at 37°C. Cultures were incubated **(A)** without perturbation (0rpm) or (**B**) with perturbation (250rpm) prior to RNA extraction (**Methods and Materials)**. Bars in each graph indicate the average value of 3 independent experiments (error bars represent standard deviation), and individual data points from biological replicates are indicated by black circles. Triangles represent a significant difference of each mutant grown in low iron conditions relative to the low iron cultures PAO1, whereas black stars indicate a significant difference in expression in high iron versus low iron conditions within a strain, as determined by one-way ANOVA with Tukey’s post-test for multiple comparisons, with a significance cutoff of p ≤ 0.05.

### PrrF-mediated AQ production is retained in static growth conditions

Our proteomics and qRT-PCR analysis indicated that AQ regulation of T6SS genes is specific to static conditions, although the underlying mechanism of this shift was not clear. Previous reports have demonstrated that static growth can dramatically impact *P. aeruginosa* physiology, including regulated expression of virulence traits. These reports have suggested that diminished oxygen availability in static conditions plays a defining role in mediating these effects (56-58). Additional studies showed that oxygen limitation decreases PQS production while increasing production of the PQS precursor molecules, AHQs (59). As such, we hypothesized that oxygen limitation resulting from static growth conditions would favor production of AHQs, potentially influencing AQ regulation of T6SS.

To test this, we used a validated LC-MS/MS method (46) to measure levels of various AQ species in shaking and static cultures of wild-type PAO1 and the isogenic Δ*prrF* mutant. As previously observed (23, 33), PrrF was required for optimal production of the C7 and C9 congeners of PQS, AQNO, and AHQ in shaking conditions (**Fig. 4A,B,C, white bars**). Similarly, PrrF was required for optimal production of all AQ congeners except HQNO in static cultures (**Fig. 4A,B,C, gray bars**), indicating that PrrF still promotes production of most AQs in static growth conditions. Production of C7-PQS, C9-PQS, and NQNO was diminished in static conditions, likely due to the previously reported oxygen requirement for activity of the PqsH and PqsL enzymes (59, 60). In contrast, production of AHQ congeners was more robust in static than in shaking conditions, both in the wild type and Δ*prrF* strains (**Fig. 4C, gray bars versus white bars**). However, when we performed analysis of cultures grown in shaking microaerobic conditions, AHQ levels were not significantly enhanced relative to shaking aerobic cultures, despite decreased production of PQS and AQNO congeners (**Fig. 4A,B,C striped bars versus white bars**). This indicates that increased AHQ levels in static cultures is not solely due to reduced oxygen availability, but that static growth causes distinct changes in *P. aeruginosa* AQ metabolite profiles.

**Figure 4.**
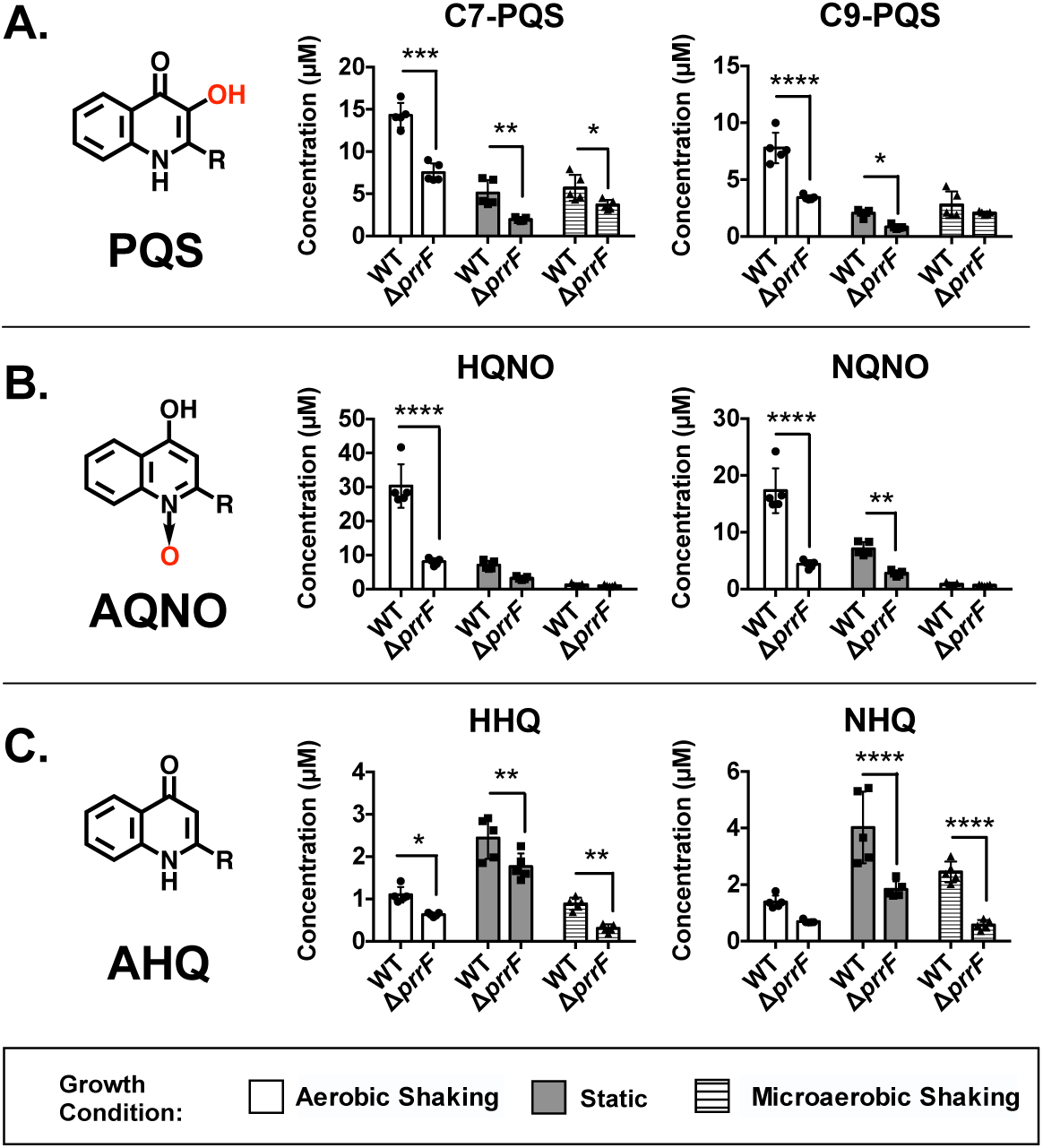
Concentrations of AQs in shaking, static, and microaerobic cultures of *P. aeruginosa.* Cultures were grown in 1.5 mL of DTSB media without iron supplementation in 14mL polystyrene round bottom culture tubes and incubated for 18 hours at 37°C. Microaerobic cultures were sealed in an air-tight GasPak™ system in the presence of an EZ Campy Container™ packet. Shaking and microaerobic cultures were incubated with perturbation (250rpm). Static cultures were incubated in shaking incubators with no perturbation (0rpm). The core structures of (**A**) PQS, (**B**) AQNO, and (**C**) AHQ molecules are depicted in black, with oxygen-dependent structural elements highlighted in red (R indicates alkyl chain). AQs were quantified by LC-MS/MS. Bars in each graph indicate the average concentration of 5 independent experiments, and individual data points from each biological replicate are indicated by black circles (error bars represent standard deviation). Asterisks indicate a significant difference compared to PAO1 within each growth condition as determined by one-way ANOVA with Sidak’s post test for multiple comparisons, with significance thresholds as follows: *p≤0.05; **p≤0.005; ***p≤0.0005; ****p≤0.00005.

### PrrF-mediated antimicrobial activity is altered by static growth conditions

We next probed the impact of altered AQ profiles in static growth on iron-regulated interactions with *S. aureus*. As discussed above, we previously discovered that iron starvation enhances AQ-dependent antimicrobial activity against *S. aureus* on agar plates as well as in a transwell co-culture system (20, 23). Surprisingly, the Δ*prrF* mutant demonstrated similar antimicrobial activity to wild type PAO1 in this previous study, suggesting that PrrF’s effects on AQ-mediated antimicrobial activity may be impacted by static growth (23). This idea was further supported by our proteomic, qPCR, and metabolite analysis discussed above, showing that both PrrF-mediated iron regulatory pathways and AQ profiles are altered in static growth (**Fig. 2 and Fig. 3)**. To investigate how culture perturbation would affect antimicrobial activity, we co-inoculated mono- and co-cultures of *P. aeruginosa* strain PAO1 and *S. aureus* strain USA300 in DTSB, supplemented with or without 100 µM FeCl_3_, and grew them either statically or in a shaking incubator at 250rpm. After 18 hours, viability of *P. aeruginosa* and *S. aureus* was quantified by enumerating CFUs on selective PIA and Baird Parker agar, respectively. In agreement with previous studies showing that AQs mediate antimicrobial activity against *S. aureus*, USA300 viability was reduced significantly during low iron co-culture with PAO1 under static conditions, whereas the Δ*pqsA* mutant did not exhibit antimicrobial activity against USA300 (**Fig. 5B, white bars**). Also in agreement with our earlier work, iron limitation significantly enhanced antimicrobial activity of PAO1 against USA300 in static co-cultures (**Fig. 5B, white versus gray bars**). Iron limitation similarly enhanced the antimicrobial activity of the Δ*prrF* mutant, although this trend was not statistically significant, perhaps due to overall weak growth of *S. aureus* in static conditions.

**Figure 5.**
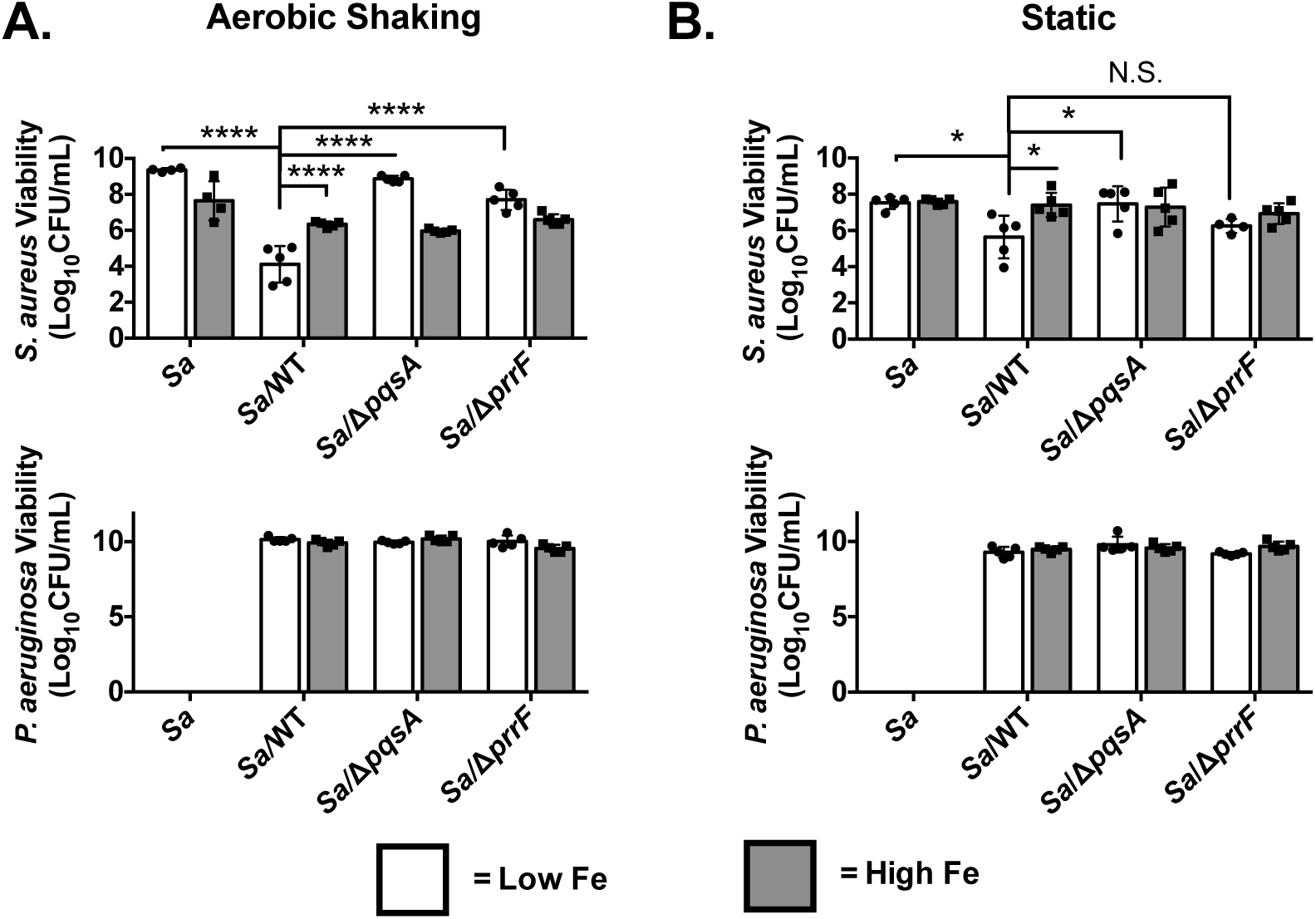
PrrF-mediated antimicrobial activity is altered in shaking conditions. *P. aeruginosa* and *S. aureus* were co-inoculated in DTSB media supplemented with (high iron) or without (low iron) 100 µM FeCl_3_. Co-cultures were incubated either in 14mL round bottom polystyrene cell culture tubes in shaking aerobic conditions **(A)** or in 6 well polystyrene cell culture plates in static conditions **(B)** for 18 hours. After incubation, CFUs were enumerated as described in the Materials and Methods. Bars in each graph indicate the average value in low (white bars) or high (gray bars) iron conditions, and individual data points from biological replicates are indicated by circles (low iron) or squares (high iron). Error bars indicate the standard deviation of 5 independent experiments. Asterisks indicate a significant difference compared to *S. aureus* monoculture as determined by one-way ANOVA with Tukey’s post test for multiple comparisons, with significance thresholds as follows: *p≤0.05; **p≤0.005; ***p≤0.0005; ****p≤0.00005.

In shaking cultures, we observed similar AQ-dependent, iron-regulated antimicrobial activity. However, *S. aureus* viability was reduced approximately 50,000-fold in low iron co-cultures with PAO1, signifying more robust killing than in static co-cultures (**Fig. 5A, white bars).** Notably, the Δ*prrF* mutant was defective for antimicrobial activity relative to the isogenic PAO1 parent strain in low iron shaking co-cultures (**Fig. 5A, white bars**), a result that contrasted with our previous and current reports on static co-cultures (23). *P. aeruginosa* viability was not affected by iron supplementation or deletion of either *pqsA* or *prrF* in any of the conditions tested (**Fig. 5**), verifying that changes in antimicrobial activity were not the result of changes in the viability of the *P. aeruginosa* Δ*pqsA* or Δ*prrF* mutants.

### Static growth alters the impact of specific AQs on antimicrobial activity

We observed that the impact of PrrF on antimicrobial activity is reduced during static growth, which was surprising since PrrF is still required for optimal production of most AQs during static growth (**Fig. 4, white and gray bars)**. Because antimicrobial activity was AQ-dependent in both shaking and static co-cultures, we next questioned whether individual AQ species differentially impacted antimicrobial activity in shaking versus static cultures. LC-MS/MS analysis showed that levels of both HQNO and NQNO are dramatically reduced in static conditions, and HQNO, which was the most abundant AQNO congener observed in our analysis of shaking cultures, was not dependent on PrrF in these conditions (**Fig. 4B**). This led us to hypothesize that AQNO-mediated antimicrobial activity would be greatly reduced, and likely unaffected by PrrF, in static conditions. To test the relative contributions of AQNOs in static versus shaking conditions, we co-incubated *S. aureus* with an AQNO-deficient PAO1 Δ*pqsL* strain in liquid co-cultures. Similar to previous analyses reported by our lab and others (20, 24), Δ*pqsL* was completely defective for antimicrobial activity against *S. aureus* in shaking cultures (**Fig. 6A).** In contrast, the Δ*pqsL* retained modest antimicrobial activity in static co-cultures (**Fig. 6B**). Considering the dramatic decrease in AQNOs during static growth, these data suggest the relative contribution of AQNOs towards antimicrobial activity is reduced in static co-cultures.

**Figure 6.**
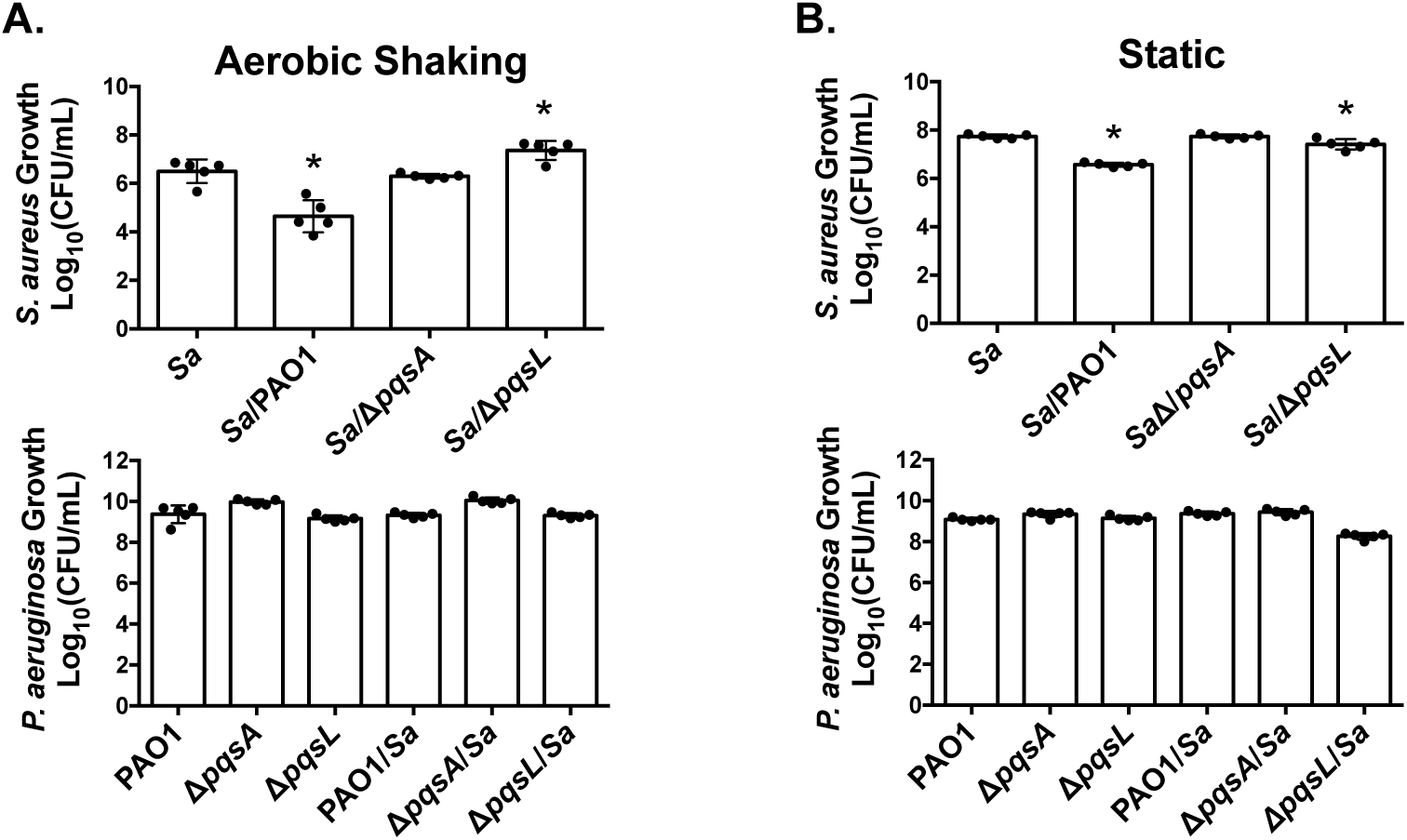
AQNO-mediated antimicrobial activity is reduced in static conditions. *P. aeruginosa* and *S. aureus* were co-inoculated in DTSB media supplemented with (high iron) or without (low iron) 100 µM FeCl_3_. Co-cultures were incubated either in 14mL round bottom polystyrene cell culture tubes in shaking aerobic conditions **(A)** or in 6 well polystyrene cell culture plates in static conditions **(B)** for 18 hours. After incubation, CFUs were enumerated as described in the **Materials and Methods**. Bars in each graph indicate the average value, and individual data points from biological replicates are indicated by circles. Error bars indicate the standard deviation of 5 independent experiments. Asterisks indicate a significant difference compared to *S. aureus* monoculture as determined by one-way ANOVA with Dunnett’s post test for multiple comparison, with a significance threshold of *p≤0.05.

Our LC-MS/MS analysis also showed that AHQs were much more abundant in static cultures compared to shaking cultures of both the wild type and Δ*prrF* mutant (**Fig. 4C, gray bars**). Thus, AHQ production by the Δ*prrF* mutant in static culture, even if it is reduced compared to wild type, may still be robust enough to promote antimicrobial activity against *S. aureus* in a seemingly PrrF-independent manner. Conversely, our LC-MS/MS analysis showed that shaking microaerobic cultures did not allow for a similar increase in AHQ levels (**Fig. 4C, striped bars**). To determine if the ability of the Δ*prrF* mutant to maintain antimicrobial activity in static growth was due solely to oxygen limitation or due to the increase in AHQs, we co-cultured *S. aureus* with *P. aeruginosa* wild type PAO1, Δ*pqsA*, or Δ*prrF* strains with shaking in either microaerobic or aerobic conditions. Similar to what was observed in shaking aerobic cultures, the Δ*prrF* mutant was defective for antimicrobial activity in shaking microaerobic conditions (**Fig. 7A,B**), indicating that changes in PrrF-dependent antimicrobial activity during static growth are not solely due to changes in oxygen availability, but may instead correlate with an overall increase in AHQ levels. Combined, these results demonstrate that different classes of AQ metabolites have distinct impacts on polymicrobial interactions in static versus shaking cultures, with AHQs potentially playing a more prominent role than AQNOs during static growth.

**Figure 7.**
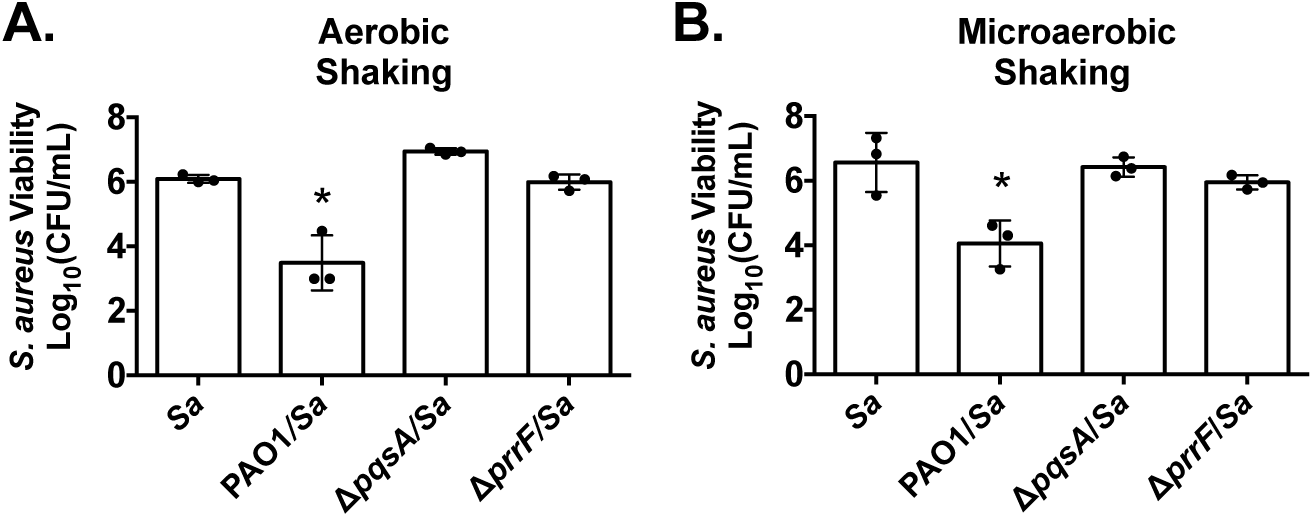
*P. aeruginosa* antimicrobial activity requirement for PrrF is not influenced by oxygen availability in co-culture with *S. aureus.* *P. aeruginosa* and *S. aureus* were co-inoculated into 1.5 mL of iron-deplete DTSB media in 14mL round-bottom polystyrene culture tubes. Cultures were incubated for 18hr at 37°C in either **(A)** shaking aerobic or in **(B)** shaking microaerobic conditions. Microaerobic cultures were incubated in air-tight GasPak systems™ in the presence of a GasPak EZ Campy Container™ packet to achieve microaerobic conditions. After 18hr incubation, CFUs were enumerated as described above. Bars in each graph indicate the average value, and individual data points from biological replicates are indicated with circles. Error bars indicate the standard deviation of 5 independent experiments. Asterisks indicate a significant difference compared to *S. aureus* monoculture as determined by one-way ANOVA with Dunnett’s post test for multiple comparisons, with a significance threshold of *p≤0.05.

## DISCUSSION

The PrrF sRNAs are critical for *P. aeruginosa* pathogenesis due to their ability to mediate numerous biological functions upon iron starvation (9). Amongst these is the Δ*prrF* mutant’s well-documented defect in the production of AQs (23, 33), which are necessary for virulence in several infection models (61-63). In the current study, we identified the HSI-II T6SS loci as a novel PrrF-responsive system, and we provide evidence that this iron regulatory effect is promoted by static growth and mediated by AQs. T6SS contributes to *P. aeruginosa* virulence in several models of infection (54, 55) and also facilitates inter-bacterial interactions in *P. aeruginosa* and other bacterial species (64, 65). T6SS mediates cell-to-cell interactions that are contact dependent and likely disrupted in shaking cultures (64), making this newly elucidated iron regulatory pathway more relevant to static growth. This study therefore provides yet another potential mechanism for how the PrrF sRNAs may contribute to pathogenesis. More broadly, our results highlight how iron regulatory pathways are highly dynamic and subject to changing environmental conditions, which may have broad implications for the heterogeneous conditions within the host.

Iron depletion was previously reported to enhance expression of T6SS systems in *P. aeruginosa*, though this was presumed to be the result of ferric uptake regulator (Fur) regulation due to the presence of a Fur consensus site in one of the promoters of the HSI-II locus (54). In the current study, we show that expression of T6SS genes is dependent on PrrF in static conditions, which themselves are regulated by iron via the Fur protein. The PrrF sRNAs appear to regulate the HSI-II locus island through AQs, likely the PQS or AHQ molecules that function as co-inducers of the PqsR/MvfR regulatory protein. Previous studies suggest that MvfR activates expression of the HSI-II T6SS loci via another quorum sensing regulator, RhlR, which directly activates HSI-II T6SS promoter activity in *P. aeruginosa* strain PAO1. Interestingly, these previous studies showed that AQs induced HSI-II T6SS in shaking cultures grown in nutrient-rich LB medium (55), whereas we found that AQs were not necessary for HSI-II T6SS expression in shaking cultures grown in a nutrient depleted medium (DTSB). There are several reasons for this apparent discrepancy, including media selection and the well-documented variability of *P. aeruginosa* strains between laboratories (66). As such, additional studies will be needed to determine the mechanism by which static growth conditions alter AQ-mediated regulation T6SS. However, it is clear that iron substantially affects T6SS gene expression, and the use of static conditions in our current study allowed us to elucidate how the PrrF sRNAs contribute to this regulatory effect.

Static growth also revealed changes in how iron affects the expression of phenazine biosynthetic proteins (**Fig. 2**). Recent studies of iron regulation in shaking conditions showed that iron starvation diminished phenazine biosynthesis gene expression and phenazine production (12, 36), while the static growth conditions in the current studies showed that iron depletion enhanced expression of phenazine biosynthesis proteins. This difference may be related to the varying role of phenazines in shaking versus static growth conditions. For instance, phenazines can induce auto-poisoning and extracellular DNA release, and these toxic effects are enhanced by nutrient depletion during aerobic shaking growth (67). However, phenazines also contribute to iron acquisition and survival due to their inherent redox-cycling activity, by which they are capable of catalyzing the formation of bioavailable ferrous iron from ferric iron (68, 69). This may be especially pertinent in static conditions, where iron requirements for biofilm formation are enhanced (70). The inherent redox-cycling activity of phenazines promotes biofilm formation by promoting the reduction of ferric iron to more bioavailable ferrous iron (68-70).

Our study also showed that pyochelin siderophore biosynthetic proteins were more strongly induced by iron depletion in static cultures as compared to shaking cultures (**Fig. 2**). While pyochelin is primarily appreciated for its role in the acquisition of ferric iron, iron-bound pyochelin can also promote the production of reactive oxygen species, particularly in conjunction to phenazines (71-74). As such, coordinated production of phenazines and pyochelin in static cultures may function to promote *P. aeruginosa* biofilm formation. Biofilms are an important adaptive feature of *P. aeruginosa* infection, and the appearance of robust biofilm-producing *P. aeruginosa* isolates during chronic lung infections is associated with increased mortality in CF patients (4, 75, 76). While the current study did not specifically address the impact of iron depletion within biofilm communities, the distinct effects of iron during static growth may be relevant to *P. aeruginosa’s* ability to form biofilms during chronic infections.

In addition to promoting virulence in mammalian hosts, many of the iron-regulated systems discussed in this report have been implicated in polymicrobial interactions. Previous work from our groups showed that iron depletion enhanced the production of AQs, thereby enhancing antimicrobial activity during co-culture with *S. aureus* (23). While this earlier work suggested that PrrF was not required for antimicrobial activity, the current study demonstrates that the role of PrrF in antimicrobial activity is dependent on culture agitation. Specifically, we show that PrrF is required for antimicrobial activity in shaking cultures, but dispensable in static cultures. Our work further highlighted distinct roles of different AQ species in mediating antimicrobial activity in shaking versus static conditions. Notably, we observed greatly diminished levels of AQNOs in static cultures, while AHQ levels were substantially increased. Thus the mechanism of antimicrobial activity against *S. aureus* is likely quite different between shaking and static cultures. Interestingly, Filkins *et al* previously showed that AQNOs were critical for antimicrobial activity against *S. aureus* in biofilms grown on CF bronchial epithelial cells (24), which might be expected to more closely mimic our static versus shaking conditions. This apparent inconsistency could be due to any number of differences between the culture conditions used here and in the Filkins study, including the time to form a mature biofilm, media selection, and the presence of mammalian cell lines. The potential role of AHQs in mediating antimicrobial activity in static cultures has yet to be elucidated but may be linked to enhanced QS-based regulation of antimicrobial factors. As such, further studies are needed to identify the extent and mechanism of AHQ activity in static cultures.

Iron availability plays a critical role in the ability of *P. aeruginosa* to express a variety of virulence traits, including the production of secondary metabolites, toxin secretion, and biofilm formation. The Fur protein and PrrF sRNAs maintain tight control over the expression of many of these activities in response to iron availability. However, these regulatory networks have not been fully characterized in static communities, which are likely to more closely reflect conditions in the host. The current study demonstrates that static growth allows for distinct iron regulatory pathways in *P. aeruginosa*, and highlights to need to further investigate iron regulation in static microbial communities such as biofilms. Such studies are likely to reveal many more striking physiological adaptations that are critical for chronic *P. aeruginosa* infections, yielding novel targets for the development of effective antimicrobial therapies.

## ACKNOWLEDGMENTS

This work was funded in part by the National Institutes of Health (NIH) grants R01 AI123320 (to AGO), T32 GM066706 (to LKB), the Cystic Fibrosis Foundation grant CFFOGLESB17G0 (to AGO), and the University of Maryland, School of Pharmacy Mass Spectrometry Center (SOP1841-IQB2014). We thank members of the Oglesby laboratory for editing of the manuscript, and Dr. Angela Wilks for thoughtful discussions and feedback as the study progressed.

## Notes

### Competing Interest Statement

The authors have declared no competing interest.

### Summary of Updates

We have updated this revised manuscript to simplify its presentation, according to previous comments from eLife. The manuscript shows that PrrF sRNA regulation of this system is mediated via alkyl-quinolones, and we provide evidence that this regulatory pathway is enhanced in static conditions. Our manuscript additionally presents how static growth alters AQ profiles in both oxygen-dependent and oxygen-independent manners, as well as how these changes impact antimicrobial activity.

## REFERENCES

1. Koch C (2002) Early infection and progression of cystic fibrosis lung disease. Pediatr Pulmonol 34(3):232-236.10.1002/ppul.10135

2. Li Z, et al. (2005) Longitudinal development of mucoid Pseudomonas aeruginosa infection and lung disease progression in children with cystic fibrosis. JAMA 293(5):581-588.10.1001/jama.293.5.581

3. Paganin P, et al. (2015) Changes in cystic fibrosis airway microbial community associated with a severe decline in lung function. PLoS One 10(4):e0124348.10.1371/journal.pone.0124348

4. Kerem E, Corey M, Gold R, & Levison H (1990) Pulmonary function and clinical course in patients with cystic fibrosis after pulmonary colonization with *Pseudomonas aeruginosa*. J Pediatr 116(5):714–719

5. Scheetz MH, et al. (2009) Morbidity associated with *Pseudomonas aeruginosa* bloodstream infections. Diagn Microbiol Infect Dis 64(3):311-319.10.1016/j.diagmicrobio.2009.02.006

6. Wisplinghoff H, et al. (2004) Nosocomial bloodstream infections in US hospitals: analysis of 24,179 cases from a prospective nationwide surveillance study. Clin Infect Dis 39(3):309-317.10.1086/421946

7. CDC (2019) Antibiotic Resistance Threats in the United States, 2019 (Atlanta, GA.), U.S. Dept of Health and Human Services.

8. Reid DW, Withers NJ, Francis L, Wilson JW, & Kotsimbos TC (2002) Iron deficiency in cystic fibrosis: relationship to lung disease severity and chronic *Pseudomonas aeruginosa* infection. Chest 121(1):48–54

9. Reinhart AA, et al. (2017) The *Pseudomonas aeruginosa* PrrF small RNAs regulate iron homeostasis during acute murine lung infection. Infection and immunity 85(5).10.1128/IAI.00764-16

10. Hood MI & Skaar EP (2012) Nutritional immunity: transition metals at the pathogen-host interface. Nat Rev Microbiol 10(8):525-537.10.1038/nrmicro2836

11. Wakeman CA, et al. (2016) The innate immune protein calprotectin promotes *Pseudomonas aeruginosa* and *Staphylococcus aureus* interaction. Nat Commun 7:11951.10.1038/ncomms11951

12. Zygiel EM, Nelson CE, Brewer LK, Oglesby-Sherrouse AG, & Nolan EM (2019) The human innate immune protein calprotectin induces iron starvation responses in Pseudomonas aeruginosa. J Biol Chem 294(10):3549-3562.10.1074/jbc.RA118.006819

13. Kim EJ, Sabra W, & Zeng AP (2003) Iron deficiency leads to inhibition of oxygen transfer and enhanced formation of virulence factors in cultures of *Pseudomonas aeruginosa* PAO1. Microbiology 149(Pt 9):2627-2634.10.1099/mic.0.26276-0

14. Vasil ML & Ochsner UA (1999) The response of *Pseudomonas aeruginosa* to iron: genetics, biochemistry and virulence. Molecular microbiology 34(3):399–413

15. Wiens JR, Vasil AI, Schurr MJ, & Vasil ML (2014) Iron-regulated expression of alginate production, mucoid phenotype, and biofilm formation by *Pseudomonas aeruginosa*. MBio 5(1):e01010-01013.10.1128/mBio.01010-13

16. Oglesby AG, et al. (2008) The influence of iron on *Pseudomonas aeruginosa* physiology: a regulatory link between iron and quorum sensing. J Biol Chem 283(23):15558-15567.10.1074/jbc.M707840200

17. Torres VJ, et al. (2010) *Staphylococcus aureus fur* regulates the expression of virulence factors that contribute to the pathogenesis of pneumonia. Infection and immunity 78(4):1618-1628.10.1128/IAI.01423-09

18. Friedman DB, et al. (2006) *Staphylococcus aureus* redirects central metabolism to increase iron availability. PLoS pathogens 2(8):e87.10.1371/journal.ppat.0020087

19. Ochsner UA & Vasil ML (1996) Gene repression by the ferric uptake regulator in *Pseudomonas aeruginosa*: cycle selection of iron-regulated genes. Proceedings of the National Academy of Sciences of the United States of America 93(9):4409–4414

20. Nguyen AT, et al. (2016) Cystic Fibrosis Isolates of *Pseudomonas aeruginosa* Retain Iron-Regulated Antimicrobial Activity against *Staphylococcus aureus* through the Action of Multiple Alkylquinolones. Front Microbiol 7:1171.10.3389/fmicb.2016.01171

21. Reinhart AA, et al. (2015) The prrF-encoded small regulatory RNAs are required for iron homeostasis and virulence of Pseudomonas aeruginosa. Infect Immun 83(3):863-875.10.1128/IAI.02707-14

22. Mashburn LM, Jett AM, Akins DR, & Whiteley M (2005) *Staphylococcus aureus* serves as an iron source for *Pseudomonas aeruginosa* during *in vivo* coculture. Journal of bacteriology 187(2):554–566

23. Nguyen AT, Jones JW, Ruge MA, Kane MA, & Oglesby-Sherrouse AG (2015) Iron Depletion Enhances Production of Antimicrobials by *Pseudomonas aeruginosa*. J Bacteriol 197(14):2265-2275.10.1128/JB.00072-15

24. Filkins LM, et al. (2015) Coculture of *Staphylococcus aureus* with *Pseudomonas aeruginosa* Drives *S. aureus* towards Fermentative Metabolism and Reduced Viability in a Cystic Fibrosis Model. J Bacteriol 197(14):2252-2264.10.1128/JB.00059-15

25. Szamosvari D & Bottcher T (2017) An Unsaturated Quinolone N-Oxide of *Pseudomonas aeruginosa* Modulates Growth and Virulence of *Staphylococcus aureus*. Angew Chem Int Ed Engl 56(25):7271-7275.10.1002/anie.201702944

26. Lightbown JW & Jackson FL (1956) Inhibition of cytochrome systems of heart muscle and certain bacteria by the antagonists of dihydrostreptomycin: 2-alkyl-4-hydroxyquinoline N-oxides. Biochem J 63(1):130–137

27. Reil E, Hofle G, Draber W, & Oettmeier W (1997) Quinolones and their N-oxides as inhibitors of mitochondrial complexes I and III. Biochim Biophys Acta 1318(1-2):291-298.10.1016/s0005-2728(96)00150-8

28. Collier DN, et al. (2002) A bacterial cell to cell signal in the lungs of cystic fibrosis patients. FEMS Microbiol Lett 215(1):41–46

29. Bredenbruch F, Geffers R, Nimtz M, Buer J, & Haussler S (2006) The *Pseudomonas aeruginosa* quinolone signal (PQS) has an iron-chelating activity. Environ Microbiol 8(8):1318-1329.10.1111/j.1462-2920.2006.01025.x

30. Pesci EC, et al. (1999) Quinolone signaling in the cell-to-cell communication system of *Pseudomonas aeruginosa*. Proceedings of the National Academy of Sciences of the United States of America 96(20):11229–11234

31. Florez C, Raab JE, Cooke AC, & Schertzer JW (2017) Membrane Distribution of the Pseudomonas Quinolone Signal Modulates Outer Membrane Vesicle Production in *Pseudomonas aeruginosa*. MBio 8(4).10.1128/mBio.01034-17

32. Coleman JP, et al. (2008) *Pseudomonas aeruginosa* PqsA is an anthranilate-coenzyme A ligase. Journal of bacteriology 190(4):1247-1255.10.1128/JB.01140-07

33. Djapgne L, et al. (2018) The Pseudomonas aeruginosa PrrF1 and PrrF2 small regulatory RNAs (sRNAs) promote 2-alkyl-4-quinolone production through redundant regulation of the antR mRNA. Journal of bacteriology.10.1128/JB.00704-17

34. Beauchene NA, et al. (2015) Impact of Anaerobiosis on Expression of the Iron-Responsive Fur and RyhB Regulons. mBio 6(6):e01947-01915.10.1128/mBio.01947-15

35. Sampathkumar B, et al. (2006) Transcriptional and translational expression patterns associated with immobilized growth of Campylobacter jejuni. Microbiology 152(Pt 2):567-577.10.1099/mic.0.28405-0

36. Nelson CE, et al. (2019) Proteomic analysis of the *Pseudomonas aeruginosa* iron starvation response reveals PrrF sRNA-dependent iron regulation of twitching motility, amino acid metabolism, and zinc homeostasis proteins. J Bacteriol.10.1128/JB.00754-18

37. Erde J, Loo RR, & Loo JA (2014) Enhanced FASP (eFASP) to increase proteome coverage and sample recovery for quantitative proteomic experiments. J Proteome Res 13(4):1885-1895.10.1021/pr4010019

38. Wisniewski JR, Zougman A, Nagaraj N, & Mann M (2009) Universal sample preparation method for proteome analysis. Nat Methods 6(5):359-362.10.1038/nmeth.1322

39. Winsor GL, et al. (2016) Enhanced annotations and features for comparing thousands of Pseudomonas genomes in the Pseudomonas genome database. Nucleic Acids Res 44(D1):D646-653.10.1093/nar/gkv1227

40. Eng JK, Fischer B, Grossmann J, & Maccoss MJ (2008) A fast SEQUEST cross correlation algorithm. J Proteome Res 7(10):4598-4602.10.1021/pr800420s

41. Dorfer V, et al. (2014) MS Amanda, a universal identification algorithm optimized for high accuracy tandem mass spectra. J Proteome Res 13(8):3679-3684.10.1021/pr500202e

42. Kall L, Canterbury JD, Weston J, Noble WS, & MacCoss MJ (2007) Semi-supervised learning for peptide identification from shotgun proteomics datasets. Nat Methods 4(11):923-925.10.1038/nmeth1113

43. Kanehisa M, Sato Y, Kawashima M, Furumichi M, & Tanabe M (2016) KEGG as a reference resource for gene and protein annotation. Nucleic Acids Res 44(D1):D457-462.10.1093/nar/gkv1070

44. Huang W, et al. (2018) PAMDB: a comprehensive Pseudomonas aeruginosa metabolome database. Nucleic Acids Res 46(D1):D575-D580.10.1093/nar/gkx1061

45. Szklarczyk D, et al. (2017) The STRING database in 2017: quality-controlled protein-protein association networks, made broadly accessible. Nucleic Acids Res 45(D1):D362-D368.10.1093/nar/gkw937

46. Brewer LK, et al. (2019) Development and Bioanalytical Method Validation of an LC-MS/MS Assay for Simultaneous Quantitation of 2-Alkyl-4(1H)-Quinolones for Application in Bacterial Cell Culture and Lung Tissue. Submitted

47. Ochsner UA, Wilderman PJ, Vasil AI, & Vasil ML (2002) GeneChip expression analysis of the iron starvation response in *Pseudomonas aeruginosa*: identification of novel pyoverdine biosynthesis genes. Mol Microbiol 45(5):1277-1287.10.1046/j.1365-2958.2002.03084.x

48. Poole K, Heinrichs DE, & Neshat S (1993) Cloning and sequence analysis of an EnvCD homologue in *Pseudomonas aeruginosa*: regulation by iron and possible involvement in the secretion of the siderophore pyoverdine. Molecular microbiology 10(3):529–544

49. Ankenbauer RG & Quan HN (1994) FptA, the Fe(III)-pyochelin receptor of *Pseudomonas aeruginosa*: a phenolate siderophore receptor homologous to hydroxamate siderophore receptors. J Bacteriol 176(2):307–319

50. Dean CR & Poole K (1993) *Cloning and characterizati*on of the ferric enterobactin receptor gene (pfeA) of Pseudomonas aeruginosa. Journal of bacteriology 175(2):317–324

51. Serino L, et al. (1997) Biosynthesis of pyochelin and dihydroaeruginoic acid requires the iron-regulated pchDCBA operon in *Pseudomonas aeruginosa*. J Bacteriol 179(1):248–257

52. Hassett DJ, et al. (1997) An operon containing fumC and sodA encoding fumarase C and manganese superoxide dismutase is controlled by the ferric uptake regulator in *Pseudomonas aeruginosa*: fur mutants produce elevated alginate levels. J Bacteriol 179(5):1452–1459

53. Wilderman PJ, et al. (2004) Identification of tandem duplicate regulatory small RNAs in *Pseudomonas aeruginosa* involved in iron homeostasis. Proceedings of the National Academy of Sciences of the United States of America 101(26):9792-9797.10.1073/pnas.0403423101

54. Sana TG, et al. (2012) The second type VI secretion system of *Pseudomonas aeruginosa* strain PAO1 is regulated by quorum sensing and Fur and modulates internalization in epithelial cells. J Biol Chem 287(32):27095-27105.10.1074/jbc.M112.376368

55. Lesic B, Starkey M, He J, Hazan R, & Rahme LG (2009) Quorum sensing differentially regulates *Pseudomonas aeruginosa* type VI secretion locus I and homologous loci II and III, which are required for pathogenesis. Microbiology 155(Pt 9):2845-2855.10.1099/mic.0.029082-0

56. Gaines JM, Carty NL, Colmer-Hamood JA, & Hamood AN (2005) Effect of static growth and different levels of environmental oxygen on toxA and ptxR expression in the *Pseudomonas aeruginosa* strain PAO1. Microbiology 151(Pt 7):2263-2275.10.1099/mic.0.27754-0

57. Wyckoff TJ, Thomas B, Hassett DJ, & Wozniak DJ (2002) Static growth of mucoid *Pseudomonas aeruginosa* selects for non-mucoid variants that have acquired flagellum-dependent motility. Microbiology 148(Pt 11):3423-3430.10.1099/00221287-148-11-3423

58. Dotsch A, et al. (2012) The *Pseudomonas aeruginosa* transcriptome in planktonic cultures and static biofilms using RNA sequencing. PLoS One 7(2):e31092.10.1371/journal.pone.0031092

59. Schertzer JW, Brown SA, & Whiteley M (2010) Oxygen levels rapidly modulate *Pseudomonas aeruginosa* social behaviours via substrate limitation of PqsH. Mol Microbiol 77(6):1527-1538.10.1111/j.1365-2958.2010.07303.x

60. Drees SL, et al. (2018) PqsL uses reduced flavin to produce 2-hydroxylaminobenzoylacetate, a preferred PqsBC substrate in alkyl quinolone biosynthesis in *Pseudomonas aeruginosa*. J Biol Chem 293(24):9345-9357.10.1074/jbc.RA117.000789

61. Deziel E, et al. (2005) The contribution of MvfR to Pseudomonas aeruginosa pathogenesis and quorum sensing circuitry regulation: multiple quorum sensing-egulated genes are modulated without affecting lasRI, rhlRI or the production of N-acyl-L-homoserine lactones. Mol Microbiol 55(4):998-1014.10.1111/j.1365-2958.2004.04448.x

62. Xiao G, et al. (2006) MvfR, a key Pseudomonas aeruginosa pathogenicity LTTR-class regulatory protein, has dual ligands. Mol Microbiol 62(6):1689-1699.10.1111/j.1365-2958.2006.05462.x

63. Gallagher LA, McKnight SL, Kuznetsova MS, Pesci EC, & Manoil C (2002) Functions required for extracellular quinolone signaling by Pseudomonas aeruginosa. J Bacteriol 184(23):6472–6480

64. Russell AB, et al. (2013) Diverse type VI secretion phospholipases are functionally plastic antibacterial effectors. Nature 496(7446):508-512.10.1038/nature12074

65. Hood RD, et al. (2010) A type VI secretion system of *Pseudomonas aeruginosa* targets a toxin to bacteria. Cell Host Microbe 7(1):25-37.10.1016/j.chom.2009.12.007

66. Chandler CE, et al. (2019) Genomic and Phenotypic Diversity among Ten Laboratory Isolates of Pseudomonas aeruginosa PAO1. J Bacteriol 201(5).10.1128/JB.00595-18

67. Meirelles LA & Newman DK (2018) Both toxic and beneficial effects of pyocyanin contribute to the lifecycle of *Pseudomonas aeruginosa*. Mol Microbiol 110(6):995-1010.10.1111/mmi.14132

68. Glasser NR, Kern SE, & Newman DK (2014) Phenazine redox cycling enhances anaerobic survival in *Pseudomonas aeruginosa* by facilitating generation of ATP and a proton-motive force. Mol Microbiol 92(2):399-412.10.1111/mmi.12566

69. Wang Y, et al. (2011) Phenazine-1-carboxylic acid promotes bacterial biofilm development via ferrous iron acquisition. J Bacteriol 193(14):3606-3617.10.1128/JB.00396-11

70. O’May CY, Sanderson K, Roddam LF, Kirov SM, & Reid DW (2009) Iron-binding compounds impair *Pseudomonas aeruginosa* biofilm formation, especially under anaerobic conditions. J Med Microbiol 58(Pt 6):765-773.10.1099/jmm.0.004416-0

71. Wang Y & Newman DK (2008) Redox reactions of phenazine antibiotics with ferric (hydr)oxides and molecular oxygen. Environ Sci Technol 42(7):2380-2386.10.1021/es702290a

72. Britigan BE, et al. (1992) Interaction of the *Pseudomonas aeruginosa* secretory products pyocyanin and pyochelin generates hydroxyl radical and causes synergistic damage to endothelial cells. Implications for *Pseudomonas-*associated tissue injury. J Clin Invest 90(6):2187-2196.10.1172/JCI116104

73. Britigan BE, Rasmussen GT, & Cox CD (1997) Augmentation of oxidant injury to human pulmonary epithelial cells by the *Pseudomonas aeruginosa* siderophore pyochelin. Infect Immun 65(3):1071–1076

74. Coffman TJ, Cox CD, Edeker BL, & Britigan BE (1990) Possible role of bacterial siderophores in inflammation. Iron bound to the *Pseudomonas* siderophore pyochelin can function as a hydroxyl radical catalyst. J Clin Invest 86(4):1030-1037.10.1172/JCI114805

75. Henry RL, Mellis CM, & Petrovic L (1992) Mucoid *Pseudomonas aeruginosa* is a marker of poor survival in cystic fibrosis. Pediatr Pulmonol 12(3):158–161

76. Pedersen SS, Hoiby N, Espersen F, & Koch C (1992) Role of alginate in infection with mucoid *Pseudomonas aeruginosa* in cystic fibrosis. Thorax 47(1):6–13

