## Supplementary Tables and Figures for "Static growth alters PrrF- and 2-alkyl-4(1*H*)-quinolone regulation of virulence trait expression in *Pseudomonas aeruginosa*"

Running title: Static growth alters *P. aeruginosa* iron regulation

Luke K. Brewer[1], Weiliang Huang[1], Brandy Hackert[1], Maureen A. Kane[1], Amanda G. Oglesby[1,2]*

University of Maryland, Baltimore, [1]School of Pharmacy, Department of Pharmaceutical Sciences and [2]School of Medicine, Department of Microbiology and Immunology, Baltimore, Maryland, 21201

**Contents:**

Supplementary Tables S1-S7
Supplementary Figures S1-S2
Supplementary References

### SUPPLEMENTARY TABLES

#### Table S1. Bacterial strains used in mono- and co-culture experiments.

| Strain | Description | Reference |
|---|---|---|
| *P. aeruginosa* | | |
| PAO1 | Wild Type laboratory strain | (1) |
| PA14 | Wild Type laboratory strain | (2) |
| Δ*prrF1,2* | *prrF1,2* deletion in PAO1 background (PrrF deficient) | (3) |
| Δ*pqsA* | *pqsA* deletion in PAO1 background (AQ deficient) | (4) |
| Δ*pqsL* | *pqsL* deletion in PAO1 background (AQNO deficient) | (5) |
| Δ*phzA1-A2* | Deletion of Phz1 and Phz2 operons in PA14 background (Phenazine deficient) | (6) |
| Δ*pchEF* | Deletion of pyochelin synthesis genes *pchE* and *pchF* (pyochelin deficient) | (7) |
| Δ*pvdA* | *pvdA* deletion in PAO1 background (pyoverdine deficient) | (8) |
| Δ*pvdA*Δ*pchEF* | *pvdA* and *pchEF* deletion in PAO1 background (pyoverdine and pyochelin deficient) | M. Vasil |
| mPAO1 | Wild Type transposon mutant parental strain | (9) |
| $\Delta pqsH_{1:Tn}$ | *pqsH* transposon insertion mutant (PQS-deficient) in mPAO1 background | (9) |
| $\Delta pqsH_{2:Tn}$ | *pqsH* transposon insertion mutant (PQS-deficient) in mPAO1 background | (9) |
| $\Delta clpV2_{A:Tn}$ | ClpV2 transposon insertion (T6SS-HSII deficient) in mPAO1 background | (9) |
| $\Delta clpV2_{B:Tn}$ | ClpV2 transposon insertion (T6SS-HSII deficient) in mPAO1 background | (9) |
| *S. aureus* | | |
| USA300 | Wild Type laboratory isolate of *S. aureus* | (10) |
| M2 | Methicillin-resistant isolate of *S. aureus* | (11) |

#### Table S2. Primer and probe sets used in qualitative RT-PCR gene expression studies.

| Gene | Forward | Reverse | Probe |
|---|---|---|---|
| *antA* | CGCCACCCTCGACTACAG | GGGCATCTCGCTGAAGAG | TCTCCTTCGCCAACGGCCAC |
| *antR* | AAACGCCTGGGCGTAGAGTT | GCAAGGTCCTGGAGGAGATGTT | ATCCATCTCCGGATCGAAGCGCA |
| *clpV2* | CTGGGCCGAATACAAGAA | GTAGATACCGTGGCTTTCC | ATGAGCCGAGCGTTGCCGAG |
| *dotU2* | GGCAACGAAAGCGAATG | GTCGAGCAACTGGAAGAA | CGCCGAAGGTCTCGTTATGGAAGC |
| *fha2* | CTACCACCAGTCGCTATG | GTGTAGAGGGTGGAAATCA | ATTGGAGCGAGCGTCTGTTGGC |
| *hsiB2* | TGCCGTTGAAGCTACTG | CGTCGAAGGTCATCTTGT | CAAGGTGGAGGACCGCAAGCC |
| *hsiC2* | GACCTGCTGGACGATTT | AGTAGTTGGCGATGATGG | ACAAGCACATCTACACCGCCGAAT |
| *icmF2* | GGAGTGATGGTGTCGATT | CAGGGTCTGCTGGATTT | CGACGACGAGATGGCGCTGGATA |
| *lip2* | GTCAATCCCGACCTCAAT | GTTCGTAGAGGCTGAAGAA | AGTGCGGCTGTTCGAGCTGAAA |
| *oprF* | GCGTTCGCAACATGAAGAAC | CTTCTTGTTGCCGGTTTCGTA | CGGTGAGTACCATGACGTTCGTGGC |
| *omlA* | ACGCAGGACATGATAGACCAGTT | TCGTTGAAGAACAGGCTGACG | CCCGACGCCAAGTGCGGTTT |
| *prrF1,2* | AACTGGTCGCGAGATCAGC | CCGTGATTAGCCTGATGAGGAG | CCCACGCAGTCGGACTCTTCAGATT |
| *sfa2* | TGCTGCTCGACGAAATC | ATCGACCTTGTGCGTATC | CGTGTTGCAGGAAGGCGAGATTCG |

**Table S3. Number of proteins differentially altered by iron in either shaking or static conditions.**

| Shaking | | | | Static | | | |
|---|---|---|---|---|---|---|---|
| Iron Induced | PrrF dependent | Iron Repressed | PrrF dependent | Iron Induced | PrrF dependent | Iron Repressed | PrrF dependent |
| 126 | 64 | 93 | 34 | 78 | 40 | 129 | 70 |

**Table S4. Proteins exhibiting PrrF-independent iron repression in static conditions.**

| Accession Number | Name | Description | WT Hi/Lo Fe LFC | | Δ*prrF* Hi/Lo Fe LFC | |
|---|---|---|---|---|---|---|
| PA0122 | RahU | putative hemolysin | -1.76 | * | -1.04 | * |
| PA0462 | | hypothetical protein | -1.06 | * | -1.49 | * |
| PA0581 | PlsY | conserved hypothetical protein | -9.97 | * | -9.97 | * |
| PA0917 | Kup | potassium uptake protein | -9.97 | * | -9.97 | * |
| PA0981 | | hypothetical protein | -9.97 | * | -9.97 | * |
| PA1000 | PqsE | Quinolone signal response protein | -1.10 | * | -1.42 | * |
| PA1150 | Pys2 | pyocin S2 | -2.40 | * | -9.97 | * |
| PA1247 | AprE | alkaline protease secretion protein | -9.97 | * | -9.97 | * |
| PA1512 | Hcp | secreted protein Hcp | -3.11 | * | -2.18 | * |
| PA1657 | HsiB2 | Type VI secretion protein | -2.18 | * | -1.55 | * |
| PA1658 | HsiC2 | Type VI secretion protein | -2.22 | * | -1.72 | * |
| PA1904 | PhzF2 | probable phenazine biosynthesis protein | -1.13 | * | -2.27 | * |
| PA1905 | PhzG2 | probable pyridoxamine 5'-phosphate oxidase | -1.11 | * | -1.35 | * |
| PA2069 | | probable carbamoyl transferase | -1.71 | * | -1.55 | * |
| PA2204 | | probable ABC transporter binding component | -1.69 | * | -3.01 | * |
| PA2300 | ChiC | chitinase | -1.29 | * | -1.03 | * |
| PA2331 | | hypothetical protein | -1.30 | * | -1.54 | * |
| PA2385 | PvdQ | 3-oxo-C12-homoserine lactone acylase | -1.83 | * | -1.86 | * |
| PA2390 | PvdT | ABC transporter-like protein | -2.79 | * | -2.58 | * |
| PA2462 | | hypothetical protein | -1.33 | * | -1.22 | * |
| PA2463 | | hypothetical protein | -1.26 | * | -1.50 | * |
| PA2719 | | hypothetical protein | -1.52 | * | -9.97 | * |
| PA2794 | | pseudaminidase | -9.97 | * | -9.97 | * |
| PA3021 | | hypothetical protein | -1.59 | * | -1.58 | * |
| PA3082 | Gbt | glycine betaine transmethylase | -1.18 | * | -1.33 | * |
| PA3186 | OprB | Glucose/carbohydrate outer membrane porin | -1.80 | * | -1.10 | * |
| PA3224 | | hypothetical protein | -9.97 | * | -9.97 | * |
| PA3227 | PpiA | peptidyl-prolyl cis-trans isomerase A | -1.29 | * | -1.51 | * |
| PA3274 | | hypothetical protein | -3.88 | * | -9.97 | * |
| PA3336 | | probable MFS transporter | -1.38 | * | -4.13 | * |
| PA3361 | LecB | fucose-binding lectin PA-IIL | -1.69 | * | -1.89 | * |
| PA3470 | | hypothetical protein | -1.66 | * | -2.75 | * |
| PA3601 | RpmE2 | conserved hypothetical protein | -3.34 | * | -2.78 | * |
| PA3865.1 | | immunity protein | -1.00 | * | -1.81 | * |
| PA4065 | | hypothetical protein | -1.75 | * | -1.16 | * |
| PA4141 | | hypothetical protein | -3.20 | * | -2.45 | * |
| PA4174 | | probable transcriptional regulator | -1.15 | * | -1.48 | * |
| PA4214 | PhzE1 | phenazine biosynthesis protein | -1.02 | * | -1.54 | * |
| PA4220 | FptB | hypothetical protein | -1.63 | * | -2.07 | * |
| PA4224 | PchG | pyochelin biosynthetic protein | -2.09 | * | -2.13 | * |
| PA4225 | PchF | pyochelin synthetase | -2.28 | * | -2.15 | * |
| PA4229 | PchC | pyochelin biosynthetic protein | -1.20 | * | -1.79 | * |
| PA4231 | PchA | salicylate biosynthesis isochorismate synthase | -1.84 | * | -2.21 | * |
| PA4242 | RpmJ | 50S ribosomal protein L36 | -2.34 | * | -3.13 | * |

| | | | | | | |
|---|---|---|---|---|---|---|
| PA4590 | Pra | protein activator | -9.97 | * | -1.70 | * |
| PA4675 | ChtA | TonB-dependent siderophore receptor | -1.97 | * | -1.70 | * |
| PA4687 | HitA | ferric iron-binding periplasmic protein HitA | -1.15 | * | -1.79 | * |
| PA4737 | | hypothetical protein | -9.97 | * | -9.97 | * |
| PA4792 | | conserved hypothetical protein | -2.25 | * | -1.39 | * |
| PA4798 | | hypothetical protein | -9.97 | * | -9.97 | * |
| PA5330 | | hypothetical protein | -1.46 | * | -1.82 | * |
| PA5447 | WbpZ | glycosyltransferase | -9.97 | * | -9.97 | * |
| PA5481 | | hypothetical protein | -3.16 | * | -2.23 | * |

Asterisks indicate FDR-adjusted significance values when comparing high iron (100μM $FeCl_3$) to low iron (0μM $FeCl_3$) cultures. Genes exhibiting both LFC values ≥1 or ≤-1 as well as a minimum FDR adjusted p-value threshold of p≤ 0.05 were considered iron regulated.

**Table S5. Proteins exhibiting PrrF-dependent iron repression in static conditions.**

| Accession Number | Name | Description | WT Hi/Lo Fe LFC | | ∆*prrF* Hi/Lo Fe LFC | |
|---|---|---|---|---|---|---|
| PA0012 | | hypothetical protein | -1.55 | * | 0.08 | |
| PA0051 | PhzH | potential phenazine-modifying enzyme | -1.49 | * | -0.94 | * |
| PA0083 | TssB1 | type VI secretion protein | -1.22 | * | -0.33 | |
| PA0090 | ClpV1 | type VI secretion protein | -1.01 | * | -0.47 | |
| PA0315 | | hypothetical protein | -1.44 | * | -0.76 | * |
| PA0391 | | hypothetical protein | -1.99 | * | -0.07 | |
| PA0454 | | conserved hypothetical protein | -1.49 | * | -1.11 | |
| PA0460 | | hypothetical protein | -1.08 | * | -0.84 | * |
| PA0483 | | probable acetyltransferase | -9.97 | * | 0.40 | |
| PA0572 | | hypothetical protein | -1.63 | * | -0.51 | |
| PA0781 | | hypothetical protein | -1.30 | * | -0.91 | * |
| PA0788 | | hypothetical protein | -9.97 | * | 0.31 | |
| PA0798 | PmtA | phospholipid methyltransferase | -1.35 | * | 0.17 | |
| PA0938 | Wzz2 | chain length determinant protein | -1.20 | * | -0.74 | * |
| PA1056 | ShaC | putative monovalent/H+ antiporter subunit D | -9.97 | * | 0.08 | |
| PA1166 | | hypothetical protein | -1.51 | * | 0.08 | |
| PA1240 | | probable enoyl-CoA hydratase/isomerase | -9.97 | * | -0.42 | |
| PA1245 | AprX | | -2.45 | * | -0.09 | |
| PA1249 | AprA | alkaline metalloproteinase precursor | -1.56 | * | -0.23 | |
| PA1311 | PhnX | 2-phosphonoacetaldehyde hydrolase | -9.97 | * | -0.23 | |
| PA1324 | | hypothetical protein | -1.30 | * | -0.55 | |
| PA1354 | | hypothetical protein | -9.97 | * | -0.76 | |
| PA1404 | | hypothetical protein | -9.97 | * | 9.97 | * |
| PA1639 | | hypothetical protein | -1.27 | * | -0.67 | |
| PA1662 | ClpV2 | type VI secretion protein | -1.56 | * | -0.91 | * |
| PA1665 | Fha2 | type VI secretion protein | -1.80 | * | -0.51 | |
| PA1666 | Lip2 | type VI secretion protein | -1.70 | * | -0.92 | |
| PA2108 | | probable decarboxylase | -1.39 | * | 0.01 | |
| PA2127 | CgrA | cupA gene regulator A | -1.44 | * | -1.08 | |
| PA2151 | GlgE | conserved hypothetical protein | -1.25 | * | -0.16 | |
| PA2163 | | hypothetical protein | -9.97 | * | -0.50 | |
| PA2290 | Gcd | glucose dehydrogenase | -1.10 | * | -0.87 | * |
| PA2365 | HsiB3 | type VI secretion protein | -1.36 | * | -0.75 | |
| PA2366 | HsiC3 | type VI secretion protein | -2.69 | * | -0.56 | |
| PA2384 | | hypothetical protein | -1.34 | * | -0.16 | |
| PA2406 | | hypothetical protein | -2.23 | * | -0.95 | |
| PA2411 | | probable thioesterase | -1.36 | * | -0.88 | * |
| PA2540 | | conserved hypothetical protein | -1.39 | * | -0.66 | |
| PA2592 | | probable periplasmic spermidine/putrescine-binding protein | -1.34 | * | 0.26 | |
| PA2774 | Tse4 | Type VI secretion effector protein | -1.73 | * | -0.86 | |
| PA3187 | | probable ATP-binding component of ABC transporter | -2.01 | * | -0.02 | |
| PA3190 | | probable binding protein component of ABC sugar transporter | -1.81 | * | -0.88 | * |

| | | | | | | |
|---|---|---|---|---|---|---|
| PA3460 | | probable acetyltransferase | -1.13 | * | -0.10 | |
| PA3520 | | hypothetical protein | -1.98 | * | 0.12 | |
| PA3724 | LasB | elastase | -1.09 | * | -0.05 | |
| PA3727 | | hypothetical protein | -1.45 | * | -0.69 | |
| PA3768 | | probable metallo-oxidoreductase | -1.17 | * | -0.97 | * |
| PA3785 | | conserved hypothetical protein | -1.18 | * | -0.75 | |
| PA3902 | Ivy | hypothetical protein | -1.20 | * | -0.97 | |
| PA4063 | | hypothetical protein | -1.17 | * | -0.80 | * |
| PA4064 | | probable ATP-binding component of ABC transporter | -1.43 | * | -0.94 | * |
| PA4066 | | hypothetical protein | -1.63 | * | -0.21 | |
| PA4142 | | probable secretion protein | -1.26 | * | 1.37 | * |
| PA4144 | | probable outer membrane protein precursor | -1.57 | * | -0.37 | |
| PA4156 | FvbA | | -1.57 | * | 1.40 | * |
| PA4175 | PrpL | protease IV | -2.74 | * | -0.41 | |
| PA4296 | PprB | two-component response regulator | -1.12 | * | -0.45 | |
| PA4345 | | hypothetical protein | -1.04 | * | -0.92 | |
| PA4502 | | probable binding protein component of ABC transporter | -9.97 | * | 0.70 | |
| PA4607 | | hypothetical protein | -1.31 | * | -0.13 | |
| PA4624 | CdrB | cyclic diguanylate-regulated TPS partner B | -1.25 | * | -0.12 | |
| PA4625 | CdrA | cyclic diguanylate-regulated TPS partner A | -1.15 | * | -0.06 | |
| PA5061 | | conserved hypothetical protein | -1.00 | * | -0.67 | |
| PA5112 | EstA | esterase | -1.56 | * | -0.92 | * |
| PA5359 | | hypothetical protein | -1.63 | * | -0.20 | |
| PA5397 | | hypothetical protein | -4.21 | * | 4.71 | * |
| PA5531 | TonB1 | | -1.04 | * | -0.49 | |

Asterisks indicate FDR-adjusted significance values when comparing high iron (100μM $FeCl_3$) to low iron (0μM $FeCl_3$) cultures. Genes exhibiting both LFC values ≥1 or ≤-1 as well as a minimum FDR adjusted p-value threshold of p≤ 0.05 were considered iron regulated.

**Table S6. Proteins exhibiting PrrF-independent iron induction in static conditions.**

| Accession Number | Name | Description | WT Hi/Lo Fe LFC | | Δ*prrF* Hi/Lo Fe LFC | |
|---|---|---|---|---|---|---|
| PA0106 | CoxA | cytochrome c oxidase, subunit I | 2.41 | * | 1.52 | * |
| PA0268 | | probable transcriptional regulator | 9.97 | * | 9.97 | * |
| PA1164 | | conserved hypothetical protein | 3.35 | * | 9.97 | * |
| PA1195 | | hypothetical protein | 9.97 | * | 9.97 | * |
| PA1486 | | beta-peptidyl aminopeptidase | 9.97 | * | 9.97 | * |
| PA1631 | | probable acyl-CoA dehydrogenase | 9.97 | * | 9.97 | * |
| PA1864 | | probable transcriptional regulator | 1.92 | * | 9.97 | * |
| PA2507 | CatA | catechol 1,2-dioxygenase | 2.51 | * | 1.18 | * |
| PA2508 | CatC | muconolactone delta-isomerase | 4.43 | * | 2.18 | * |
| PA2512 | AntA | anthranilate dioxygenase large subunit | 2.07 | * | 1.27 | * |
| PA2513 | AntB | anthranilate dioxygenase small subunit | 3.39 | * | 2.43 | * |
| PA2514 | AntC | anthranilate dioxygenase reductase | 2.97 | * | 2.46 | * |
| PA2682 | | conserved hypothetical protein | 2.05 | * | 1.70 | * |
| PA2940 | | probable acyl-CoA thiolase | 9.97 | * | 9.97 | * |
| PA3027 | | probable transcriptional regulator | 9.97 | * | 9.97 | * |
| PA3091 | | hypothetical protein | 1.88 | * | 1.35 | * |
| PA3113 | TrpF | N-(5'phosphoribosyl)anthranilate (PRA) isomerase | 9.97 | * | 9.97 | * |
| PA3238 | | hypothetical protein | 9.97 | * | 9.97 | * |
| PA3591 | | probable enoyl-CoA hydratase/isomerase | 9.97 | * | 9.97 | * |
| PA3602 | | conserved hypothetical protein | 1.32 | * | 1.08 | * |
| PA3671 | | probable permease of ABC transporter | 1.21 | * | 1.64 | * |
| PA3672 | | Prob. ATP-binding component of ABC transporter | 1.31 | * | 1.36 | * |
| PA4030 | | conserved hypothetical protein | 1.21 | * | 1.77 | * |
| PA4154 | | conserved hypothetical protein | 1.13 | * | 1.47 | * |
| PA4635 | | conserved hypothetical protein | 1.04 | * | 2.29 | * |
| PA4957 | Psd | phosphatidylserine decarboxylase | 1.03 | * | 1.10 | * |
| PA4973 | ThiC | thiamin biosynthesis protein ThiC | 1.40 | * | 1.98 | * |
| PA4977 | AruI | 2-ketoaroxylase, AruI | 9.97 | * | 9.97 | * |
| PA5202 | | hypothetical protein | 1.88 | * | 1.69 | * |
| PA5360 | PhoB | two-component response regulator PhoB | 1.44 | * | 2.37 | * |
| PA5365 | PhoU | phosphate uptake regulatory protein | 1.17 | * | 1.74 | * |
| PA5367 | PstA | membrane protein component of ABC phosphate transporter | 1.35 | * | 1.77 | * |
| PA5368 | PstC | membrane protein component of ABC phosphate transporter | 1.65 | * | 2.25 | * |
| PA5421 | FdhA | glutathione-independent formaldehyde dehydrogenase | 1.53 | * | 1.15 | * |
| PA5437 | | probable transcriptional regulator | 9.97 | * | 9.97 | * |

Asterisks indicate FDR-adjusted significance values when comparing high iron (100μM $FeCl_3$) to low iron (0μM $FeCl_3$) cultures. Genes exhibiting both LFC values ≥1 or ≤-1 and a minimum FDR adjusted p-value threshold of p≤ 0.05 were considered iron regulated.

**Table S7. Proteins exhibiting PrrF-dependent iron induction in static conditions.**

| Accession Number | Name | Description | WT Hi/Lo Fe LFC | | ∆*prrF* Hi/Lo Fe LFC | |
|---|---|---|---|---|---|---|
| PA0044 | ExoT | exoenzyme T | 9.97 | * | 0.42 | |
| PA0091 | VgrG1 | type VI secretion system protein | 9.97 | * | 0.56 | |
| PA0127 | | hypothetical protein | 1.36 | * | 0.25 | |
| PA0297 | SpuA | probable glutamine amidotransferase | 1.21 | * | -0.33 | |
| PA0352 | | probable transporter | 1.22 | * | 0.03 | |
| PA0554 | | hypothetical protein | 1.46 | * | 0.44 | |
| PA0574 | | hypothetical protein | 9.97 | * | 0.24 | |
| PA0846 | CysZ | probable sulfate uptake protein | 9.97 | * | -0.31 | |
| PA1035 | | hypothetical protein | 1.44 | * | -1.09 | * |
| PA1062 | | hypothetical protein | 1.45 | * | -1.17 | |
| PA1148 | Eta | exotoxin A precursor | 9.97 | * | -2.65 | * |
| PA1310 | PhnW | 2-aminoethylphosphonate:pyruvate aminotransferase | 9.97 | * | -0.48 | |
| PA1606 | | hypothetical protein | 1.67 | * | 0.92 | |
| PA1814 | KerV | hypothetical protein | 2.34 | * | 0.35 | |
| PA1819 | | probable amino acid permease | 9.97 | * | -0.67 | |
| PA1959 | Upk | bacitracin resistance protein | 9.97 | * | -0.59 | |
| PA2118 | Ada | O6-methylguanine-DNA methyltransferase | 9.97 | * | -9.97 | * |
| PA2136 | | hypothetical protein | 1.53 | * | -9.97 | * |
| PA2264 | | conserved hypothetical protein | 1.16 | * | 0.54 | |
| PA2266 | | probable cytochrome c precursor | 1.07 | * | 0.60 | |
| PA2344 | MtlZ | fructokinase | 1.74 | * | -0.79 | |
| PA2509 | CatB | muconate cycloisomerase I | 1.72 | * | 0.24 | |
| PA2577 | | probable transcriptional regulator | 9.97 | * | -9.97 | * |
| PA3000 | AroP1 | aromatic amino acid transport protein | 9.97 | * | -0.36 | |
| PA3121 | LeuC | 3-isopropylmalate dehydratase large subunit | 1.29 | * | 0.56 | |
| PA3669 | | hypothetical protein | 1.03 | * | 0.90 | * |
| PA3670 | | hypothetical protein | 1.15 | * | 0.86 | * |
| PA3693 | | conserved hypothetical protein | 1.51 | * | 0.19 | |
| PA4024 | EutB | ethanolamine ammonia-lyase large subunit | 9.97 | * | -9.97 | * |
| PA4124 | HpcB | homoprotocatechuate 2,3-dioxygenase | 9.97 | * | 0.82 | |
| PA4130 | | probable sulfite or nitrite reductase | 1.23 | * | 0.33 | |
| PA4286 | | hypothetical protein | 1.06 | * | 0.06 | |
| PA4599 | MexC | RND multidrug efflux membrane fusion protein | 1.50 | * | 0.51 | |
| PA4621 | | probable oxidoreductase | 1.14 | * | 0.60 | |
| PA4734 | | hypothetical protein | 1.69 | * | 0.12 | |
| PA4835 | | hypothetical protein | 9.97 | * | -9.97 | * |
| PA4844 | CtpL | methyl accepting chemotaxis protein | 1.10 | * | 0.01 | |
| PA5185 | | conserved hypothetical protein | 9.97 | * | 0.06 | |
| PA5415 | GlyA1 | serine hydroxymethyltransferase | 1.40 | * | 0.82 | |
| PA5502 | | hypothetical protein | 1.07 | * | 0.96 | * |

Asterisks indicate FDR-adjusted significance values when comparing high iron (100μM $FeCl_3$) to low iron (0μM $FeCl_3$) cultures. Genes exhibiting both LFC values ≥1 or ≤-1 and a minimum FDR adjusted p-value threshold of p≤ 0.05 were considered iron regulated.

### SUPPLEMENTARY FIGURES

| | Whole Culture | | Supernatant (% of whole culture) | |
|---|---|---|---|---|
| | Shaking | Static | Shaking | Static |
| C7-PQS | 24.38μM ± 3.4μM | 18.34μM ± 3.58μM* | 9.66μM ± 0.954μM (39.62% ± 3.17%) | 9.49μM ± 0.57μM (51.73% ± 11.04%)^ |
| C9-PQS | 28.82μM ± 3.91μM | 14.31μM ± 2.80μM* | 2.78μM ± 0.43μM (9.63% ± 0.45%) | 4.07μM ± 0.44μM* (28.47% ± 3.85%)^ |
| HHQ | 1.50μM ± 0.20μM | 5.65μM ± 0.82μM* | 1.23μM ± 0.07μM (82.48% ± 9.54%) | 3.93μM ± 0.21μM* (69.51% ± 10.31%) |
| NHQ | 3.15μM ± 0.57μM | 14.10μM ± 2.31μM* | 1.46μM ± 0.17μM (47.24% ± 4.91%) | 5.11μM ± 0.59μM* (36.21% ± 3.55%)^ |

Wild type and Δ*pqsA* cultures were grown in 1.5 mL of DTSB media supplemented with either 0μM $FeCl_3$ in 14mL polystyrene round bottom culture tubes for 18 hours at 37˚C. Cultures were incubated without perturbation (0rpm) or with perturbation (250rpm) prior to RNA extraction (Methods and Materials). 300μL of cells were either immediately subjected to ethyl acetate extraction, or were centrifuged for 4 minutes at 21000rpm and supernatants subjected to ethyl acetate extraction. Asterisks indicate significance values when comparing static to shaking values with a minimum significance thresholf of P≤0.05. Carrots indicate significance values of percent AQ exported in static conditions versus shaking conditions with a minimum significance threshold of p≤0.05.

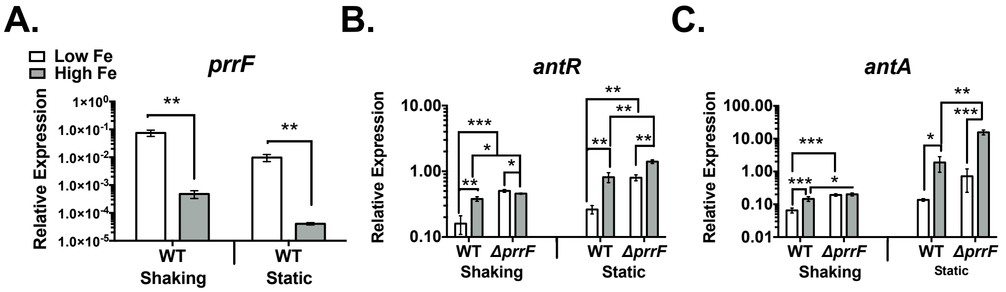


**Figure S1. PrrF expression and regulation of anthranilate catabolism genes in shaking and static conditions.** Wild type PAO1 and Δ*prrF* mutant strains were incubated in 1.5mL dialyzed trypticase soy broth (DTSB) supplemented with either low iron (white bars) or high iron (gray bars) in 14mL round bottom polystyrene culture tubes. Cultures were incubated for 18hr at 37˚C in either shaking aerobic conditions or static conditions prior to RNA extraction (**Methods and Materials).** Levels of the **(A)** *prrF*, **(B)** *antR*, and **(C)** *antA* RNAs were measured using qRT-PCR. Bars in each graph indicate the average value of 5 independent experiments (error bars represent standard deviation). Asterisks indicate a significant difference as determined by a two-tailed Student's *t* test as follows: *p<0.05; **p<0.005; ***p<0.0005.

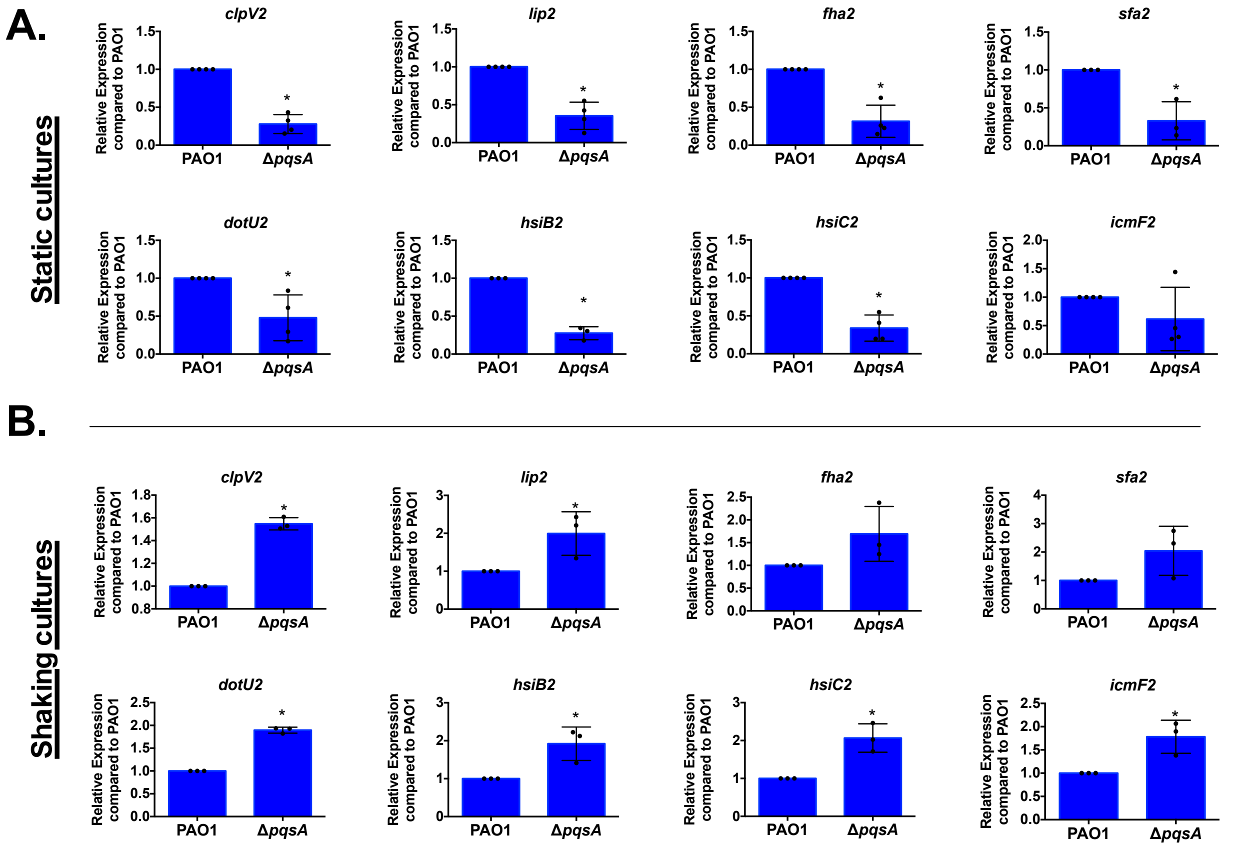


**Figure S2. Alkyl quinolones induce T6SS genes in Δ*pqsA* mutant in shaking cultures but not static cultures.** Wild type and Δ*pqsA* cultures were grown in 1.5 mL of DTSB media supplemented with either 0μM $FeCl_3$ or 100μM $FeCl_3$ in 14mL polystyrene round bottom culture tubes for 18 hours at 37˚C. Cultures were incubated without perturbation (0rpm) **(A)** or with perturbation (250rpm) **(B)** prior to RNA extraction (**Methods and Materials)**. Bars in each graph indicate the average value of 3 independent experiments (error bars represent standard deviation), and individual data points from biological replicates are indicated by circles. Aserisks represent a significant difference relative to low iron PAO1 as determined by a two-tailed Student's *t* test as follows: *p<0.05.

### Supplementary References


1. Holloway BW (1955) Genetic recombination in *Pseudomonas aeruginosa*. *J Gen Microbiol* 13(3):572-581.
2. Schroth MNC, J.J.; Green, S.K., Kominos, S.D. (1977) Epidemiology of *Pseudomonas aeruginosa* in agricultural areas. *Pseudomonas aeruginosa: Ecological Aspects and Patient Colonization*, ed Young VM (Raven Press, New York), pp 1-29.
3. Oglesby AG*, et al.* (2008) The influence of iron on *Pseudomonas aeruginosa* physiology: a regulatory link between iron and quorum sensing. *J Biol Chem* 283(23):15558-15567.
4. Nguyen AT*, et al.* (2014) Adaptation of iron homeostasis pathways by a *Pseudomonas aeruginosa* pyoverdine mutant in the cystic fibrosis lung. *J Bacteriol* 196(12):2265-2276.
5. D'Argenio DA, Calfee MW, Rainey PB, & Pesci EC (2002) Autolysis and autoaggregation in *Pseudomonas aeruginosa* colony morphology mutants. *J Bacteriol* 184(23):6481-6489.
6. Dietrich LE, Price-Whelan A, Petersen A, Whiteley M, & Newman DK (2006) The phenazine pyocyanin is a terminal signalling factor in the quorum sensing network of *Pseudomonas aeruginosa*. *Mol Microbiol* 61(5):1308-1321.
7. Banin E, Vasil ML, & Greenberg EP (2005) Iron and *Pseudomonas aeruginosa* biofilm formation. *Proc Natl Acad Sci U S A* 102(31):11076-11081.
8. Ochsner UA, Wilderman PJ, Vasil AI, & Vasil ML (2002) GeneChip expression analysis of the iron starvation response in *Pseudomonas aeruginosa*: identification of novel pyoverdine biosynthesis genes. *Mol Microbiol* 45(5):1277-1287.
9. Jacobs MA*, et al.* (2003) Comprehensive transposon mutant library of *Pseudomonas aeruginosa*. *Proc Natl Acad Sci U S A* 100(24):14339-14344.
10. Centers for Disease C & Prevention (2001) Methicillin-resistant *Staphylococcus aureus* skin or soft tissue infections in a state prison--Mississippi, 2000. *MMWR Morb Mortal Wkly Rep* 50(42):919-922.
11. Harro JM*, et al.* (2013) Draft Genome Sequence of the Methicillin-Resistant *Staphylococcus aureus* Isolate MRSA-M2. *Genome Announc* 1(1).
